# Structure, dynamics, and processing of 8oxoG:A in the nucleosome

**DOI:** 10.1101/2025.09.02.673820

**Authors:** Abigayle F. Vito, Justin A. Ling, Julia C. Ferrara, Caleb S. Jacques, Natacha Gillet, Roy González-Alemán, Yuya Qiu, Mohammad Hashemian, Carlos H. Trasvina-Arenas, Sheila S. David, Sarah Delaney, Emmanuelle Bignon, Bret D. Freudenthal

**Author notes:** To whom correspondence should be addressed; Emails: Sarah Delaney, Emmanuelle Bignon, and Bret D. Freudenthal. These authors contributed equally to this work (Abigayle F. Vito, Justin A. Ling).

## Abstract

Eukaryotic genomic DNA is packaged into chromatin through a repeating unit known as the nucleosome. In this chromatin environment, DNA is constantly exposed to several sources of DNA damage, such as reactive oxygen species (ROS), which can lead to the formation of 8-oxo-7,8-dihydroguanine (8oxoG). 8oxoG can base pair with cytosine (8oxoG:C) or form a mutagenic base pair with adenine (8oxoG:A), which can lead to single base transversions if left unrepaired. To date, the structure and dynamics of these two possible 8oxoG base pairs in the nucleosome remain unclear. Furthermore, whether MutY homologue (MUTYH) excises 8oxoG:A base pairs in the nucleosome remains elusive. Here using a combination of cryogenic-electron microscopy, molecular dynamics simulations, and biochemistry we determined the structure and dynamics of 8oxoG:C and 8oxoG:A base pairs in the nucleosome and characterize MUTYH activity in nucleosomal DNA. We found that nucleosomal 8oxoG:C forms a stable base pair using its *anti* conformation, while nucleosomal 8oxoG:A forms a more dynamic base pair using its *syn* conformation that is unable to be processed by MUTYH. This work provides fundamental insight into the accommodation of oxidative damage in the nucleosome and how this damage contributes to increased mutagenic transversions in nucleosomal compared to linker DNA.

## Introduction

In eukaryotes, genomic DNA is packaged into the nucleus in the form of chromatin which consists of a fundamental repeating unit called the nucleosome. The nucleosome is comprised of ∼147 base pairs of DNA wrapped around an octamer of histone proteins containing two copies of histones H2A, H2B, H3 and H4^1^. Both nucleosomal and non-nucleosomal DNA is susceptible to several forms of DNA damage, one of the most common being oxidative damage^2^. Oxidative damage arises from excessive levels of reactive oxygen species (ROS) which are capable of oxidizing macromolecules in the cell^3,4^. The nucleobase guanine is highly susceptible to oxidation, forming the DNA lesion 8-oxo-7,8-dihydroguanine (8oxoG)^5^. If not repaired, 8oxoG can inappropriately base pair with adenine during replication resulting in G to T (and complementary C to A) transversions, which are a common feature of cancer genomes with the single base substitution signature 18 (SBS 18)^6–10^.

8oxoG lesions are primarily repaired through the base excision repair (BER) pathway^7,11–13^. BER is initiated by DNA glycosylases that recognize and excise the modified nucleobase. The primary DNA glycosylase responsible for identifying and removing 8oxoG is 8-oxo-guanine glycosylase 1 (OGG1)^12^. OGG1 specifically recognizes 8oxoG across from dC and excises 8oxoG leaving an abasic site that is further processed by downstream BER enzymes^11,14–16^. If OGG1 does not identify and excise 8oxoG prior to DNA replication, replicative polymerases can insert dA complementary to 8oxoG forming a mutagenic base pair^6,17^. A second DNA glycosylase, MutY homologue (MUTYH), identifies and excises dA opposite 8oxoG, resulting in an abasic site that is further processed by downstream BER enzymes^18–20^. This MUTYH-initiated BER allows the potential for restoration of the 8oxoG:C base pair which can then be recognized by OGG1 to avoid imprinting a mutation into the genome.

Chromatin structure was recently shown to be a key determinant of oxidation-induced mutations in human genomes where G to T mutations tend to persist in regions of tightly packed chromatin following oxidative stress^21,22^. This mutational signature is seemingly due to lack of efficient DNA repair as oxidative lesions were formed in a relatively uniform manner across tightly packed chromatin and linker DNA regions^9,10,21,23,24^. Consistent with these observations, the compact nature of chromatin structure has been implicated in limiting the ability of DNA repair enzymes to access their DNA substrates^21,25–28^. DNA damage within the nucleosome is differentially accessible to repair enzymes depending on its position within the nucleosome and prior studies suggest that accessibility of the DNA damage site dictates repair efficiency^28–32^. Indeed, recent studies have shown that OGG1 can access 8oxoG within a nucleosome at some solvent-exposed rotational orientations^21,27,28,33–37^, however, how oxidative damage impacts chromatin structure and whether MUTYH can function on nucleosomal DNA remains elusive. Here, we address these gaps in knowledge using a combination of structural, computational, and biochemical techniques. We found that the presence of 8oxoG in a nucleosome does not impact the base pairing properties of 8oxoG or global nucleosome structure at the positions tested. Classical MD simulations reveal that the presence of 8oxoG in the nucleosome induces only very local structural perturbations to the DNA backbone and that the nucleobases in the 8oxoG:A base pair exhibit a specific dynamical behavior which is not present in the 8oxoG:C base pair. Finally, biochemical assays reveal that MUTYH is not appreciably active in the context of the nucleosome.

## Results

To investigate the effect of oxidative damage to chromatin structure, we generated nucleosomes containing 8oxoG opposite dC or dA within a 147 bp Widom 601 strong positioning sequence and investigated the molecular structure and dynamics of the nucleosome core particle (NCP). It has been previously shown that DNA-histone contacts vary greatly in stability between the dyad and entry/exit sites in a nucleosome^38,39^. This difference in stability led us to investigate whether the structure and dynamics of the 8oxoG lesion is dependent upon the translational position of the 8oxoG:C or 8oxoG:A base pair. To this end, we determined cryo-EM structures of 8oxoG:C or 8oxoG:A at four distinct superhelical locations (SHL): SHL_+2_, SHL_+3_, SHL_+4_, and SHL_−6_ (Fig. 1). These structures were used to understand the base pairing properties off 8oxoG within the nucleosome and whether the nucleosome is impacted by the presence of 8oxoG.

**Fig. 1:**
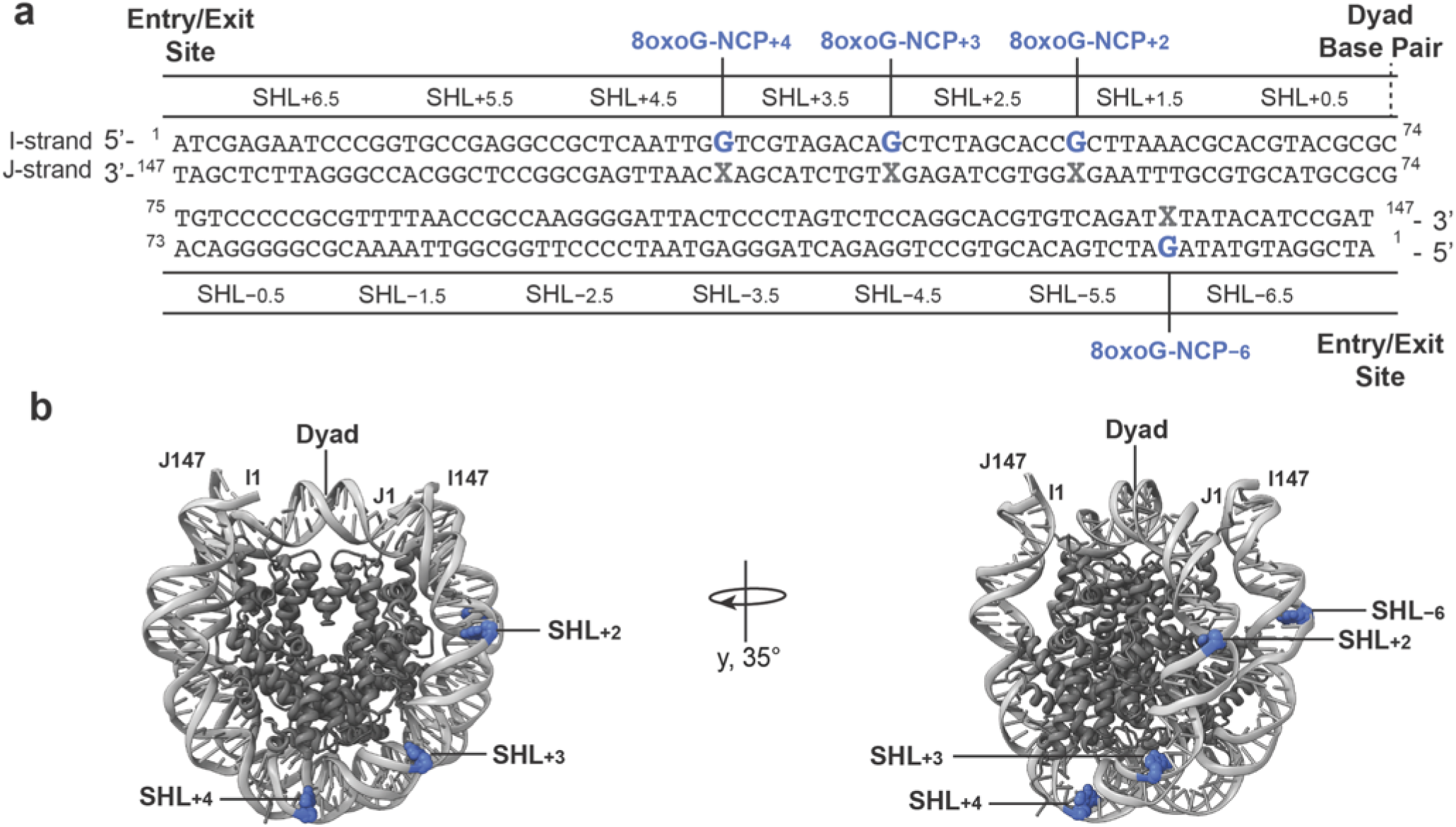
Translational positions for nucleosomal 8oxoG. **a,** Diagram of the Widom 601 strong positioning sequence highlighting the 8oxoG mutation sites used to reconstitute damaged NCPs. 8oxoG sites are shown in blue and their complementary bases (C or A) are shown in gray. **b,** Two orientations of the nucleosome core particle (PDB: 9DS4) with 8oxoG positions at SHL_+2_, SHL_+3_, SHL_+4_, and SHL_−6_ labeled.

### Structure and dynamics of a non-mutagenic 8oxoG:dC base pair in the nucleosome

To obtain insight into the structure and dynamics of the non-mutagenic 8oxoG:C base pair in the nucleosome, we generated a recombinant nucleosome containing an 8oxoG:C base pair at SHL_+2_ (8oxoG:C-NCP_+2_) (SFig. 1). At SHL_+2_, the 8oxoG:C base pair is positioned two superhelical turns away from the nucleosome dyad where the 8oxoG is solvent-exposed and the complementary dC is facing toward the histone (i.e., histone-occluded) (Fig. 1b). We performed single particle analysis and obtained a 2.9 Å cryo-EM reconstruction of 8oxoG:C-NCP_+2_ (Fig. 2a-b, SFig. 2, STable 1). Importantly, the 8oxoG:C-NCP_+2_ reconstruction was of sufficient quality to readily assign the register of the nucleosomal DNA and accurately define the position the 8oxoG:C base pair (SFig. 3). At this position, 8oxoG adopts the *anti* conformation and forms classic Watson-Crick base pairing with the complementary dC (Fig. 2c). This conformation is the same as seen in a previously determined high resolution x-ray crystal structure of an 8oxoG:C base pair in non-nucleosomal DNA^40^, indicating the nucleosome does not alter the base pairing properties of 8oxoG:C at SHL_+2_. To further understand whether the 8oxoG:C base pair at SHL_+2_ impacts overall nucleosome structure, we determined a 3.8 Å cryo-EM structure of a nucleosome without DNA damage (non-damaged, ND-NCP) (SFig.1, SFig. 4-5). Structural comparison of 8oxoG:C-NCP_+2_ and ND-NCP revealed minimal changes to the histone octamer (RMSD: 0.4 Å) and nucleosomal DNA (RMSD: 0.4 Å) (Fig. 2d), suggesting the 8oxoG:C base pair at SHL_+2_ does not impact global nucleosome structure, consistent with prior observations^41^.

**Fig. 2:**
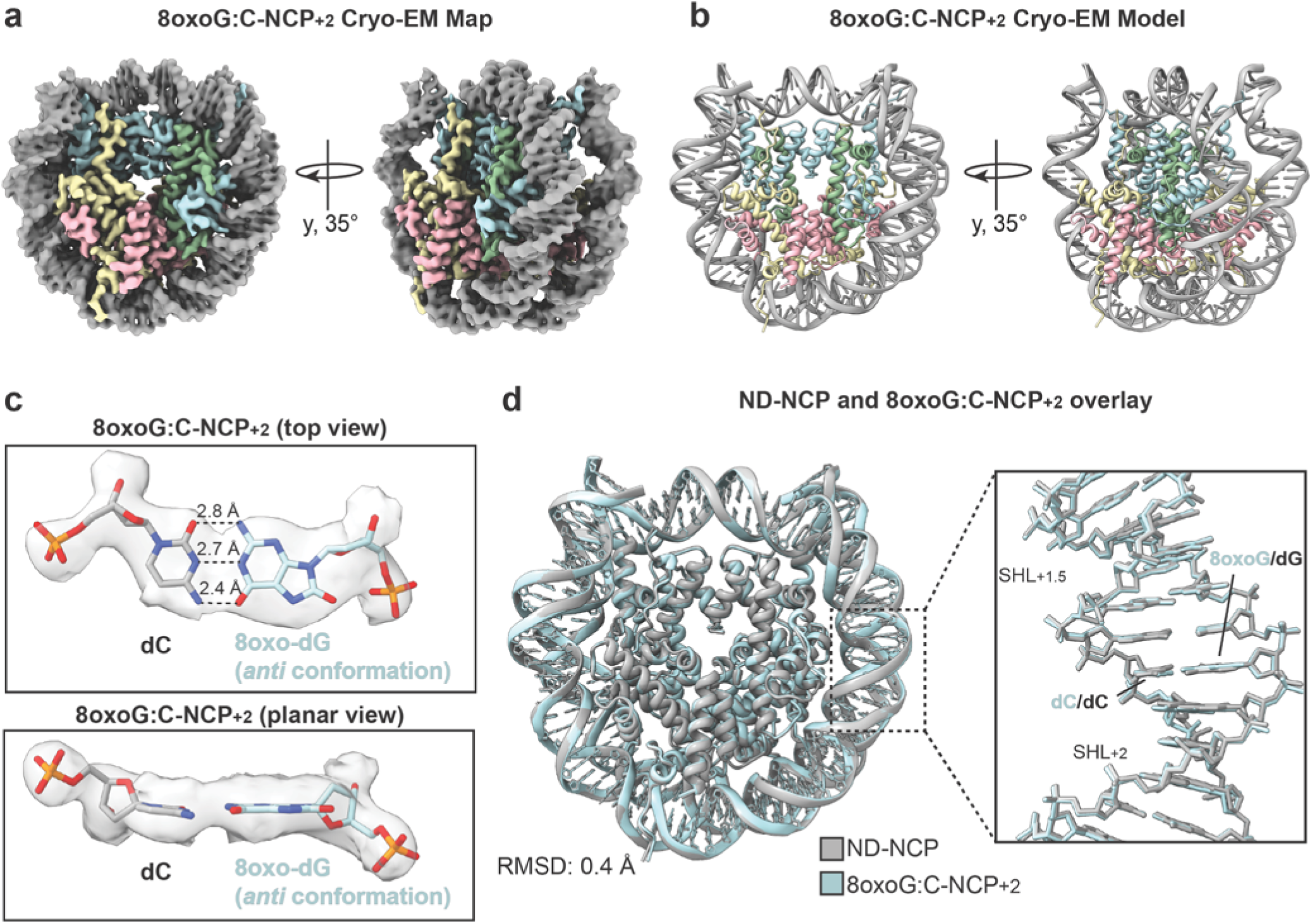
Structural determination of the 8oxoG:C base pair at SHL_+2_ in the nucleosome. **a,** Final 8oxoG:C-NCP_+2_ cryo-EM map shown in two orientations. **b,** Final 8oxoG:C-NCP_+2_ model shown in the same two orientations as (a). **c,** Top view and planar view of the 8oxoG:C base pair at SHL_+2_. The density from the cryo-EM map is shown as a transparent gray surface. The dotted lines represent hydrogen bonds between bases and are labeled with their respective lengths. **d,** Structural comparison of ND-NCP (gray) and 8oxoG:C-NCP_+2_ (blue). RMSD value represents difference between the histone octamers.

To determine if the base pairing properties of the 8oxoG:C base pair and its impact on overall nucleosome structure are position dependent, we generated a recombinant nucleosome containing an 8oxoG:C base pair at SHL_+3_ (8oxoG:C-NCP_+3_) (SFig. 1). At this superhelical position, the 8oxoG:C base pair is positioned adjacent to SHL_+2_, one helical turn further from the dyad. Like 8oxoG:C-NCP_+2_, 8oxoG:C-NCP_+3_ features a solvent-exposed 8oxoG and a histone-occluded complementary dC (Fig. 1b). We performed single particle analysis and obtained a 3.3 Å cryo-EM reconstruction of 8oxoG:C-NCP_+3_ (Fig. 3a-b, SFig. 6, STable 1). Importantly, the 8oxoG:C-NCP_+3_ reconstruction was of sufficient quality to assign the register of the DNA and correctly define the position of the 8oxoG:C base pair (SFig. 7). At this position, the 8oxoG lesion adopts the *anti* conformation and forms Watson-Crick base pairing (Fig. 3c). This conformation is the same as seen for 8oxoG:C base pairing in non-nucleosomal DNA and our 8oxoG:C-NCP_+2_ (Fig. 2c). Furthermore, we performed structural comparison of 8oxoG:C-NCP_+3_ to our ND-NCP and found minimal changes to the histone octamer (RMSD: 0.4 Å) and nucleosomal DNA (RMSD: 0.4 Å) (Fig. 3d). In addition, we previously reported two nucleosome structures containing 8oxoG:C at SHL_+4_ (8oxoG:C-NCP_+4_) and SHL_−6_ (8oxoG:C-NCP_−6_) at resolutions of 3.0 Å and 3.1 Å, respectively (Fig. 3e-f). In both structures, the 8oxoG is solvent-exposed and the complementary dC is histone-occluded^21^. At SHL_+4_, the 8oxoG:C base pair is positioned adjacent to SHL_+3_, one helical turn further from the dyad. At SHL_−6_, the 8oxoG:C base pair is near the entry exit site on the opposite face of the nucleosome (Fig 1b). Importantly, in both the 8oxoG:C-NCP_+4_ and 8oxoG:C-NCP_−6_ the 8oxoG lesion adopts the *anti* conformation and forms classic Watson-Crick base pairing with the complementary dC (Fig. 3g-h). Cumulatively, these data indicate that the position of the 8oxoG lesion in the nucleosome does not alter the base pairing conformation or base pairing properties of the non-mutagenic 8oxoG:C base pair. Furthermore, we performed structural comparison of 8oxoG:C-NCP_+4_, and 8oxoG:C-NCP_−6_ to our ND-NCP. We observed minimal structural changes to the histone octamer and nucleosomal DNA for 8oxoG:C at SHL_+4_ and SHL_−6_ indicating the 8oxoG lesion does not alter the global nucleosome structure at any position tested.

**Fig. 3:**
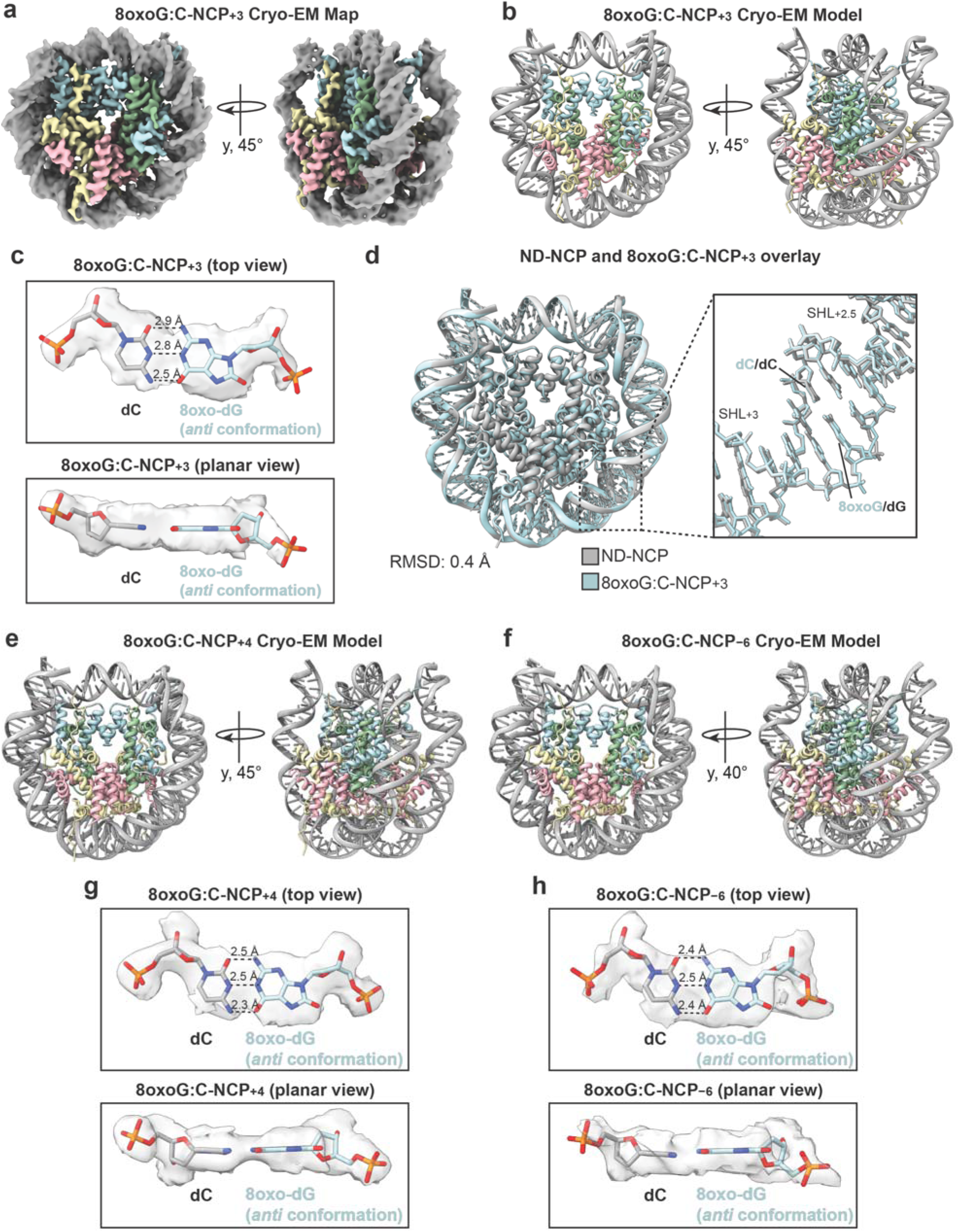
Accommodation of the 8oxoG:C base pair in the nucleosome is the same across multiple translational positions. **a,** Final 8oxoG:C-NCP_+3_ cryo-EM map shown in two orientations. **b,** Final 8oxoG:C-NCP_+3_ model shown in the same two orientations as (a). **C,** Top view and planar view of the 8oxoG:C base pair at SHL_+3_. The density from the cryo-EM map is shown as a transparent gray surface. The dotted lines represent hydrogen bonds between bases and are labeled with their respective lengths. **d,** Structural comparison of ND-NCP (gray) and 8oxoG:C-NCP_+3_ (blue). RMSD value represents difference between the histone octamers. **e,** Final model from previously reported structure of 8oxoG:C-NCP_+4_ shown in the same two orientations as (a)^21^ **f,** Final model from previously reported structure of 8oxoG:C-NCP_−6_ shown in the same two orientations as (a)^21^ **g,** Top view and planar view of the 8oxoG:C base pair at SHL_+4_. The density from the cryo-EM map is shown as a transparent gray surface. The dotted lines represent hydrogen bonds between bases and are labeled with their respective lengths. **h,** Top view and planar view of the 8oxoG:C base pair at SHL_−6_. The density from the cryo-EM map is shown as a transparent gray surface. The dotted lines represent hydrogen bonds between bases and are labeled with their respective lengths.

The 8oxoG:C-NCP cryo-EM structures represent a stable, single conformation of the 8oxoG:C base pair within the nucleosome structure. To further understand the stability and dynamics of the 8oxoG:C base pair, we performed five replicates of 1 µs classical molecular dynamics (MD) simulations on the 8oxoG:C-NCP_+2_ structure. Additionally, we performed five replicates of 1 µs classical MD simulations on the ND-NCP and three replicates of 1 µs classical MD simulations on a 21-bp non-nucleosomal (i.e. duplex) DNA oligonucleotide with and without an 8oxoG:C base pair. Analysis of the intra-base pair structural parameters reveal that the G:C and 8oxoG:C base pairs in the ND-NCP and 8oxoG:C-NCP_+2_ systems both exhibit a monotonous dynamical behavior and maintain classical Watson-Crick base pairing (Fig. 4a).

**Fig. 4:**
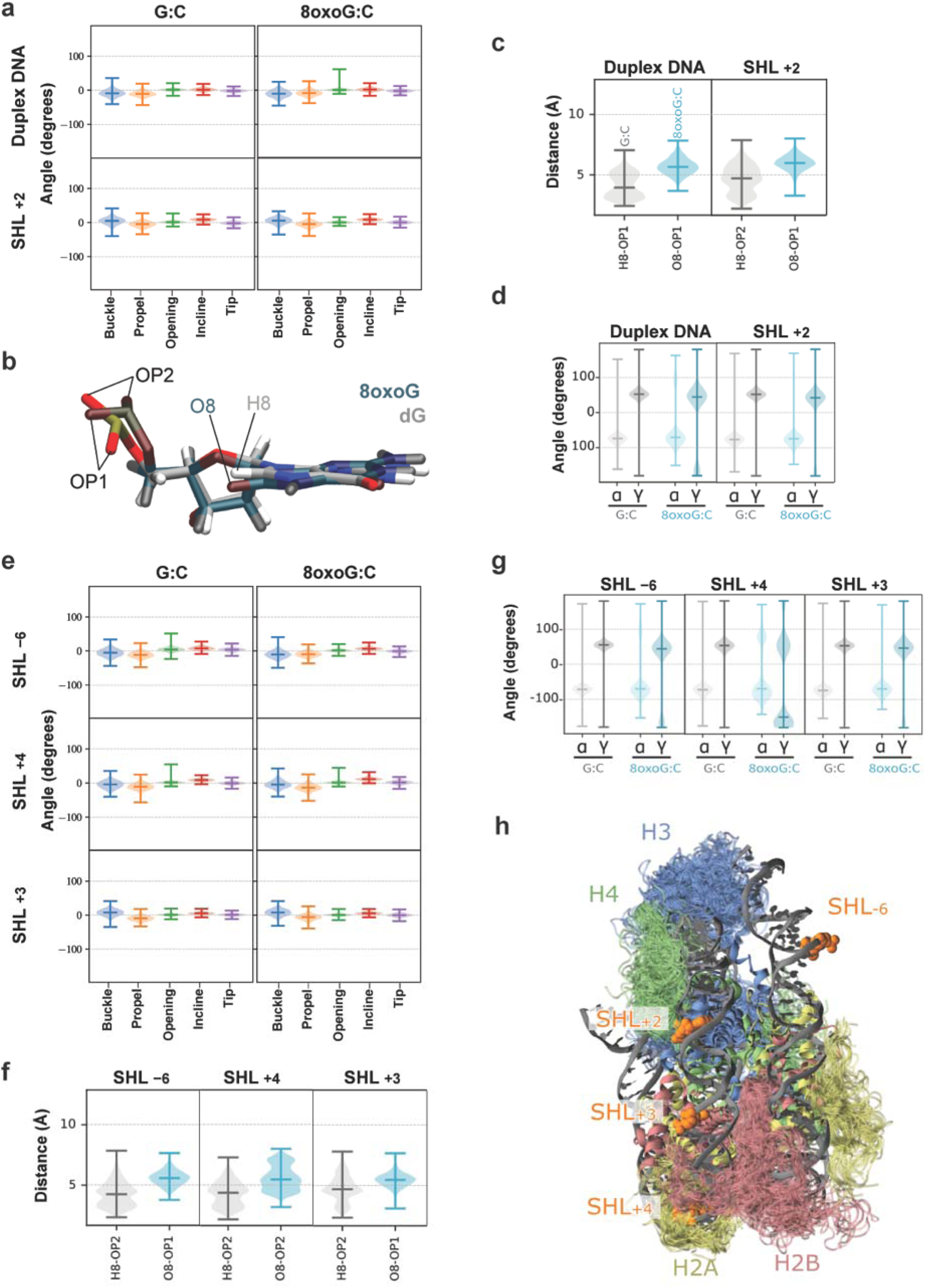
Dynamics of the 8oxoG:C base pair in the nucleosome and non-nucleosomal (duplex) DNA. **a,** Distribution of the intra-base pair angles for the control and 8oxoG:C systems in duplex DNA or at SHL_+2_. **b,** Superimposition of dG (grey) and 8oxoG (blue/green). Non-bridging oxygen atoms on the phosphate group (OP1 and OP2) and closest atom on the nucleobase (H8 for dG or O8 for 8oxoG) are labeled. **c,** Distribution of the difference between either the H8 (undamaged, grey) or O8 (for 8oxoG:C, blue) atom and the closest non-bridging oxygen atom on the phosphate group (OP1 or OP2) calculated from MD simulations in duplex DNA or SHL_+2_. **d,** Distribution of the α and γ backbone angles of 8oxoG or dG in the control, for duplex DNA or SHL_+2_. **e,** Distribution of the intra-base pair angles for the control and 8oxoG:C systems at SHL_−6_, SHL_+4_, and SHL_+3_. **f,** Distribution of the difference between either the H8 (undamaged, grey) or O8 (for 8oxoG:C, blue) atom and the closest non-bridging oxygen atom on the phosphate group (OP1 or OP2) calculated from MD simulations at SHL_−6_, SHL_+4_, and SHL_+3_. **g,** Distribution of the α and γ backbone angles of 8oxoG or dG in the control, for SHL_−6_, SHL_+4_, and SHL_+3_. **h,** Side view of the projection of histone tail conformations sampled by MD simulations. The SHL of the lesion is highlighted in orange.

This trend is consistent across non-nucleosomal DNA and nucleosomal DNA at SHL_+2_, indicating that DNA compaction in the nucleosome does not alter the Watson-Crick base pairing dynamics or stability. Additionally, in agreement with the cryo-EM structures, the MD trajectories show that the presence of the 8oxoG:C base pair does not disturb the global nucleosome structure. However, a few subtle rearrangements are observed in the local DNA backbone that allow for the accommodation of the extra O8 atom present in 8oxoG. Indeed, the distance between O8 of 8oxoG and the closest non-bridging oxygen on the phosphate (OP1 or OP2) in the 8oxoG:C-NCP_+2_ system is greater than the distance between the H8 of the reference dG and the closest non-bridging oxygen in the ND-NCP system (Fig. 4b-c), indicating that there are local rearrangements of the DNA backbone due to the presence of 8oxoG. This local rearrangement of the DNA backbone is observed in both the 8oxoG:C-NCP_+2_ and duplex DNA system. In the nucleosome as well as in non-nucleosomal DNA, the presence of the O8 atom on 8oxoG induces the formation of an organized first solvation shell around this atom, which is absent in the ND-NCP system (SFig. 8). One or two water molecules can bridge O8 of 8oxoG and the DNA backbone. Similarly, sodium cations are present within 3 Å of the O8 of 8oxoG atom (SFig. 9), and these interactions can impact the DNA backbone angle distribution. However, at SHL_+2_, we do not observe differences in the α and γ dihedral angles of the 8oxoG nucleotide compared to the control dG in the ND-NCP system (Fig. 4d).

To determine whether the intra-base pair structural parameters of 8oxoG:C and the 8oxoG-induced local rearrangements in the DNA backbone are position dependent in the nucleosome, we performed five replicates of 1 µs classical MD simulations on the 8oxoG:C-NCP_+3_, 8oxoG:C-NCP_+4_, and 8oxoG:C-NCP_−6_ structures. We found that at each additional SHL tested, the intra-base pair structural parameters follow the same monotonous dynamical behavior as seen in the 8oxoG:C-NCP_+2_ system and maintain Watson-Crick base pairing (Fig. 4e). These similarities indicate that the intra-base pair parameters and stability of the 8oxoG:C base pair are not position dependent in the nucleosome. Additionally, we compared the DNA backbone perturbation at SHL_+3_, SHL_+4_, and SHL_−6_ to SHL_+2_ to determine if the 8oxoG:C-induced local DNA backbone rearrangements are position dependent. We found that in all systems, the distance between O8 of 8oxoG and the closest non-bridging oxygen is greater than the distance between the H8 of dG and the closest non-bridging oxygen in the ND-NCP system (Fig. 4b,f), similar to what was observed at SHL_+2_. Additionally, we observed that the α and γ dihedral angles of the 8oxoG nucleotide exhibit a position-dependent bimodal distribution for 8oxoG:C (Fig. 4g). This bimodality, especially pronounced at SHL_+4_, might be a result of contacts made between the 8oxoG:C base pair and the histone H2A tail at this site (Fig. 4h). Cumulatively, these data indicate that the position of the 8oxoG:C base pair in the nucleosome does not alter the local rearrangements in the DNA backbone apart from the position-dependent α and γ dihedral angle distribution of the backbone of the 8oxoG nucleotide.

### Structure and dynamics of a mutagenic 8oxoG:dA base pair in the nucleosome

To obtain insight into the structure and dynamics of the mutagenic 8oxoG:A base pair in the nucleosome, we generated a recombinant nucleosome containing an 8oxoG:A base pair at SHL_+2_ (8oxoG:A-NCP_+2_) (SFig. 1). In this position, the 8oxoG is solvent-exposed and the complementary dA is histone-occluded. We performed single particle analysis and determined a 2.7 Å cryo-EM structure of 8oxoG:A-NCP_+2_ (Fig. 5a-b, SFig. 10, STable 2). The 8oxoG:A-NCP_+2_ reconstruction was of sufficient quality to assign the register of the nucleosomal DNA and define the position of the 8oxoG:A base pair. (SFig. 11). At SHL_+2_, 8oxoG adopts the *syn* conformation and forms Hoogsteen base pairing with the complementary dA (Fig 5c). This conformation is the same as seen in a previously determined high resolution x-ray crystal structure of an 8oxoG:A base pair in non-nucleosomal DNA^42^, indicating that nucleosome structure does not significantly alter the base pair conformation or base pairing properties of 8oxoG:A at SHL_+2_. To determine whether the presence of the 8oxoG:A base pair at SHL_+2_ impacts overall nucleosome structure, we performed structural comparison of 8oxoG:A-NCP_+2_ to our ND-NCP. This comparison revealed minimal changes to both the histone octamer (RMSD: 0.4 Å) and nucleosomal DNA (RMSD: 0.4 Å) (Fig 5d), indicating that the 8oxoG:A base pair at SHL_+2_ does not impact global nucleosome structure.

**Fig 5:**
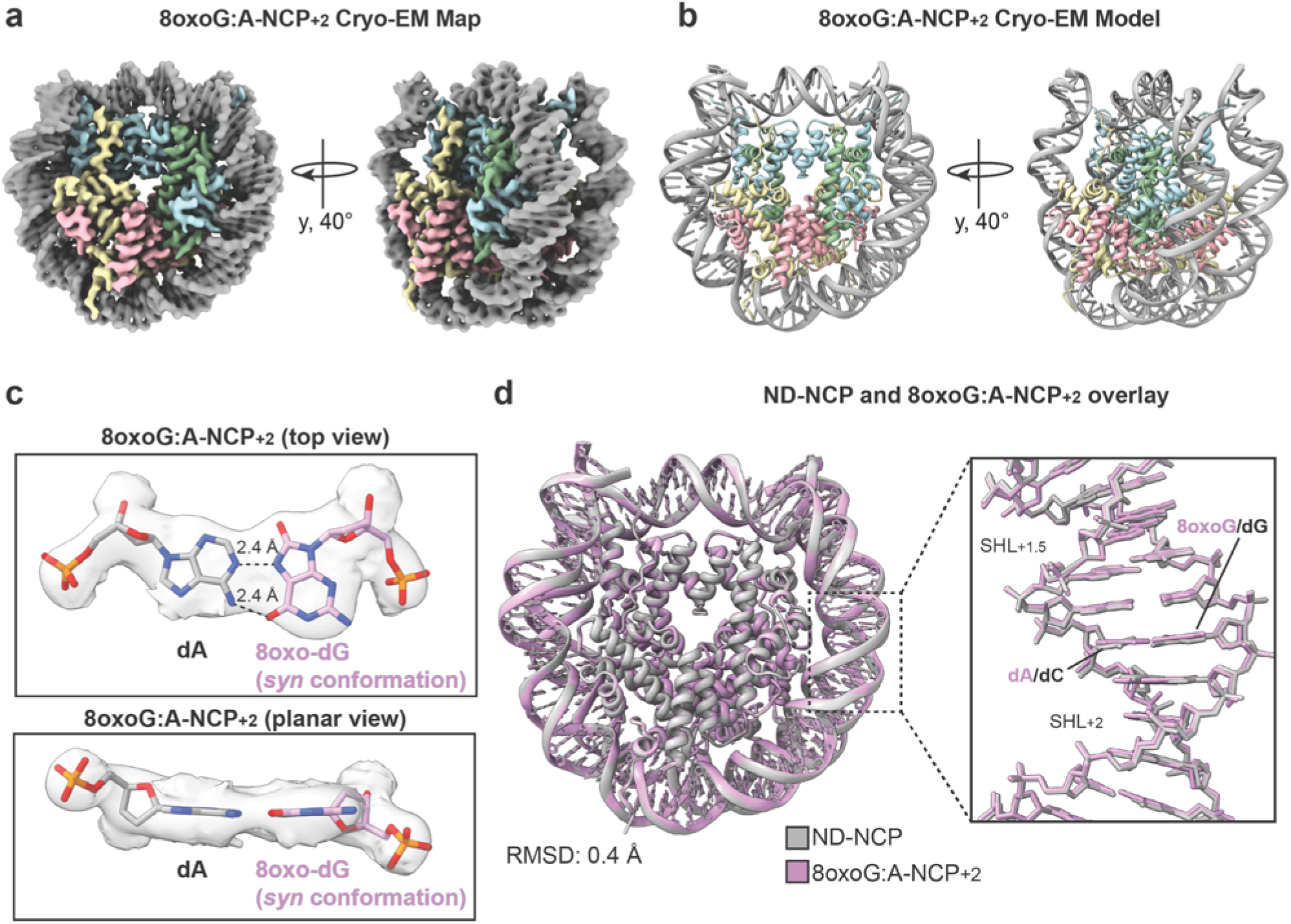
Structural determination of the 8oxoG:A base pair at SHL_+2_ in the nucleosome. **a,** Final 8oxoG:A-NCP_+2_ cryo-EM map shown in two orientations. **b,** Final 8oxoG:A-NCP_+2_ model shown in the same two orientations as (a). **c,** Top view and planar view of the 8oxoG:A base pair at SHL_+2_. The density from the cryo-EM map is shown as a transparent gray surface. The dotted lines represent hydrogen bonds between bases and are labeled with their respective lengths. **d,** Structural comparison of ND-NCP (gray) and 8oxoG:A-NCP_+2_ (pink). RMSD value represents difference between the histone octamers.

To understand if the base pairing properties of 8oxoG:A and its impact on overall nucleosome structure is position dependent, we generated three additional recombinant nucleosomes containing an 8oxoG:A base pair at SHL_+3_ (8oxoG:A-NCP_+3_), SHL_+4_ (8oxoG:A-NCP_+4_), and SHL_−6_ (8oxoG:A-NCP_−6_) (SFig. 1). These positions are consistent with those used for analysis of the non-mutagenic 8oxoG:C base pair. We then performed single particle analysis and obtained a 2.6 Å, 2.8 Å, and 2.9 Å cryo-EM reconstruction of 8oxoG:A-NCP_+3_, 8oxoG:A-NCP_+4_, and 8oxoG:A-NCP_−6_, respectively (Fig. 6, STable 2). All three reconstructions were of sufficient quality to assign the register of the nucleosomal DNA and define the position of the 8oxoG:A base pair (SFig. 12-16). In the 8oxoG:A-NCP_+3_, 8oxoG:A-NCP_+4_, and 8oxoG:A-NCP_−6_ structures, the 8oxoG lesion adopts the *syn* conformation and forms Hoogsteen base pairing with the complementary dA (Fig. 7a-c). This conformation is the same as observed for 8oxoG:A base pairing on non-nucleosomal DNA and our 8oxoG:A-NCP_+2_ (Fig 5c). Notably, these data indicate that the position of the 8oxoG lesion in the nucleosome does not alter the base pairing conformation or base pairing properties of the mutagenic 8oxoG:A base pair. To understand whether the position of the 8oxoG:A base pair in the nucleosome impacts overall nucleosome structure, we performed structural comparison of 8oxoG:A-NCP_+3_, 8oxoG:A-NCP_+4_, and 8oxoG:A-NCP_−6_, to our ND-NCP. We observed minimal structural changes to the histone octamer for 8oxoG:A-NCP_+3_ (RMSD: 0.5 Å), 8oxoG:A-NCP_+4_ (RMSD: 0.5 Å), and 8oxoG:A-NCP_−6_ (RMSD: 0.5 Å), compared to the ND-NCP (Fig. 7d-f). There were also minimal changes in the nucleosomal DNA for 8oxoG:A-NCP_+3_ (RMSD: 0.5 Å), 8oxoG:A-NCP_+4_ (RMSD: 0.5 Å), and 8oxoG:A-NCP_−6_ (RMSD: 0.5 Å), compared to the ND-NCP. Cumulatively, these data indicate the 8oxoG:A base pair does not alter global nucleosome structure at any position tested.

**Fig. 6:**
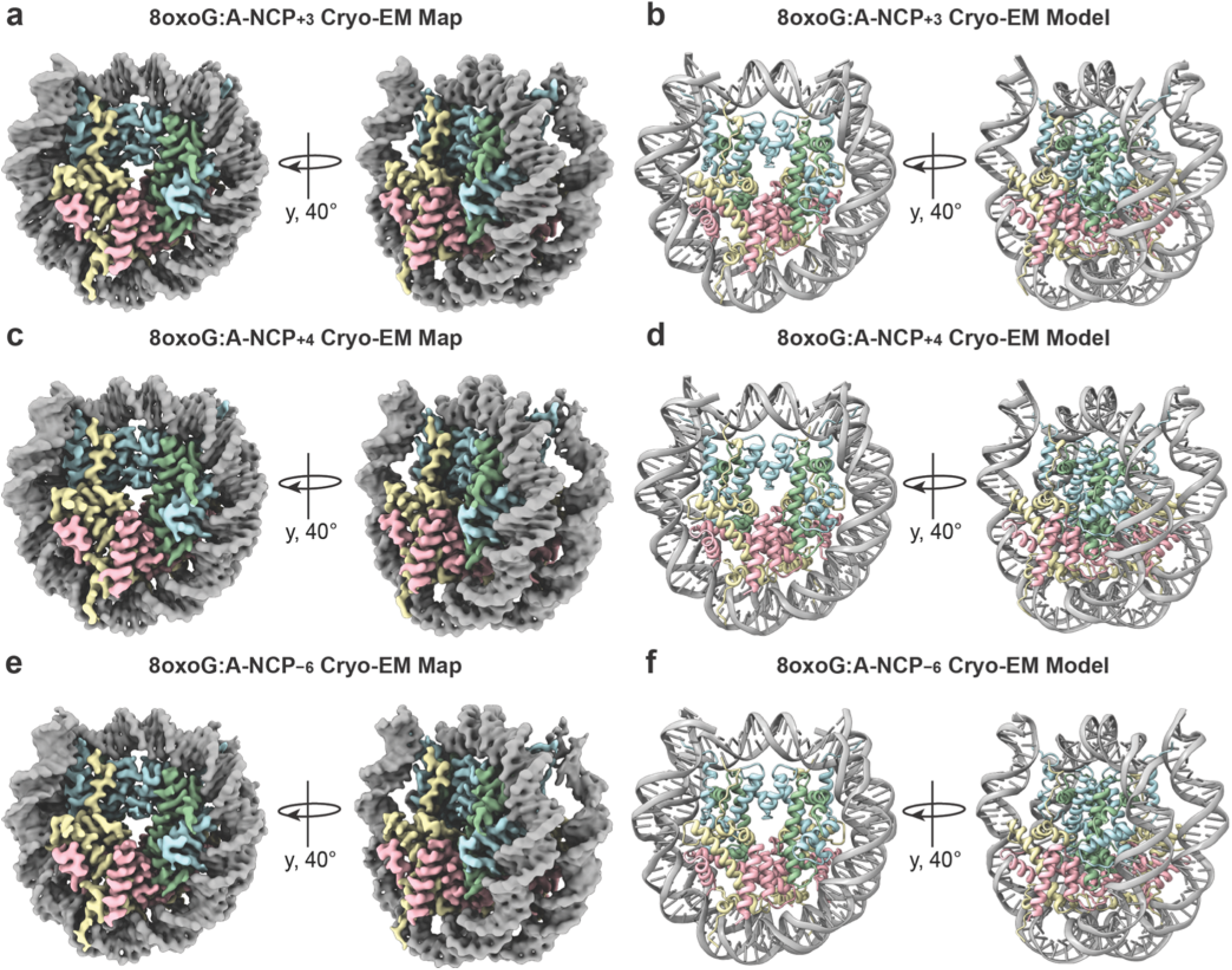
Structural determination of the 8oxoG:A base pair at multiple translational positions in the nucleosome. **a,** Final 8oxoG:A-NCP_+3_ cryo-EM map shown in two orientations. **b,** Final 8oxoG:A-NCP_+3_ model shown in the same two orientations. **c,** Final 8oxoG:A-NCP_+4_ cryo-EM map shown in two orientations. **d,** Final 8oxoG:A-NCP_+4_ model shown in the same two orientations. **e,** Final 8oxoG:A-NCP_−6_ cryo-EM map shown in two orientations. **f,** Final 8oxoG:A-NCP_−6_ model shown in the same two orientations.

**Fig. 7:**
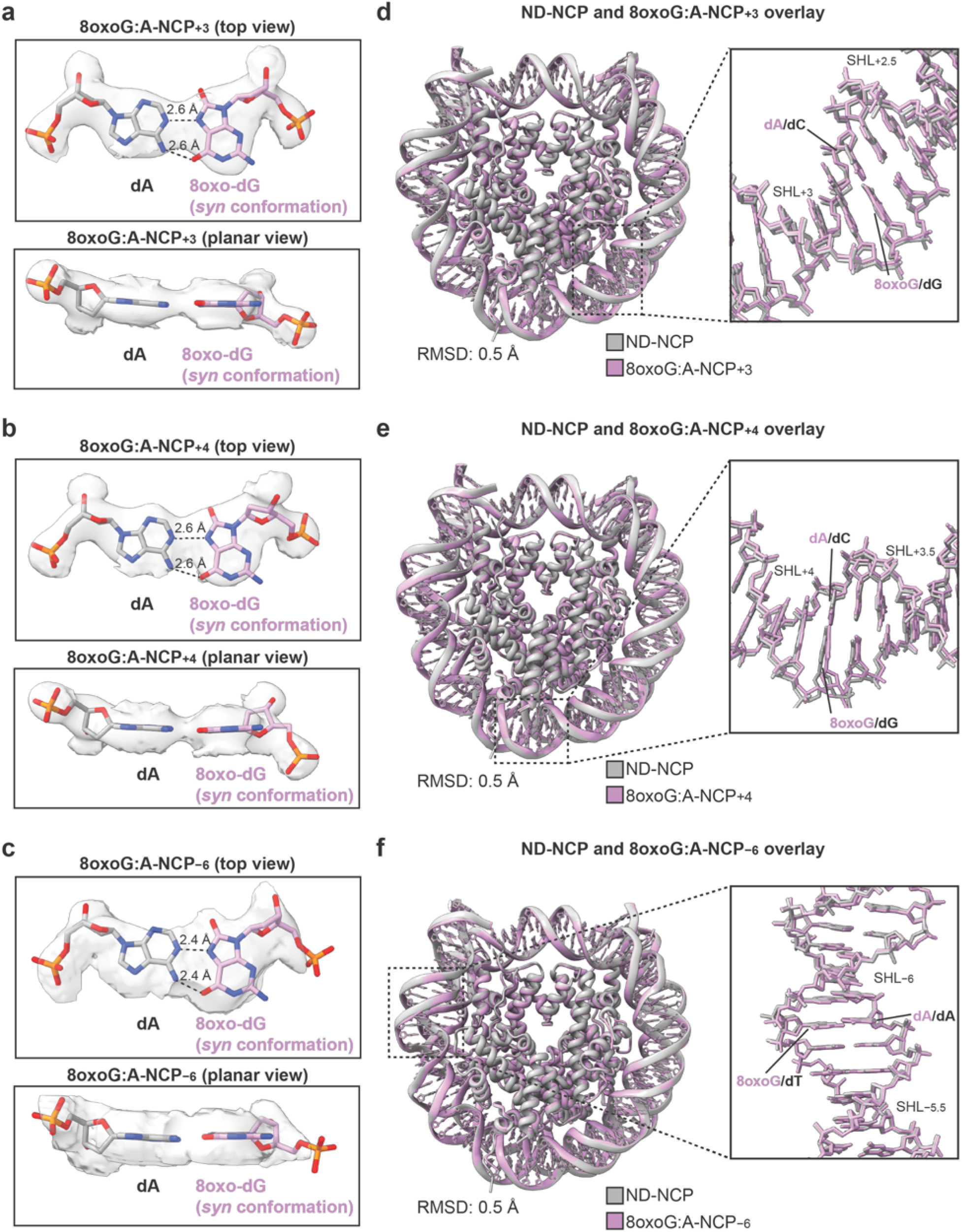
Accommodation of the 8oxoG:A base pair in the nucleosome is the same across multiple translational positions. **a,** Top view and planar view of the 8oxoG:A base pair at SHL_+3_. The density from the cryo-EM map is shown as a transparent gray surface. The dotted lines represent hydrogen bonds between bases and are labeled with their respective lengths. **b,** Structural comparison of ND-NCP (gray) and 8oxoG:A-NCP_+3_ (pink). RMSD value represents difference between the histone octamers. **c,** Top view and planar view of the 8oxoG:A base pair at SHL_+4_. The density from the cryo-EM map is shown as a transparent gray surface. The dotted lines represent hydrogen bonds between bases and are labeled with their respective lengths. **d,** Structural comparison of ND-NCP (gray) and 8oxoG:A-NCP_+4_ (pink). RMSD value represents difference between the histone octamers. **e,** Top view and planar view of the 8oxoG:A base pair at SHL_−6_. The density from the cryo-EM map is shown as a transparent gray surface. The dotted lines represent hydrogen bonds between bases and are labeled with their respective lengths. **f,** Structural comparison of ND-NCP (gray) and 8oxoG:A-NCP_−6_ (pink). RMSD value represents difference between the histone octamers.

The 8oxoG:A-NCP cryo-EM structures represent a stable, single conformation of the 8oxoG:A base pair and nucleosome structure. To further understand the stability and dynamics of the 8oxoG:A base pair, we performed five replicates of 1 µs MD simulations on the 8oxoG:A-NCP_+2_ structure and three replicates of 1 µs MD simulations on a 21-bp non-nucleosomal DNA oligonucleotide with and without an 8oxoG:A base pair. We found that the mismatch Hoogsteen base pairing exhibits a specific dynamical behavior within the DNA helix. Analysis of the intra-base pair parameters reveals that, while the 8oxoG:C Watson-Crick base pairing exhibits a monotonous dynamical behavior (Fig. 4a), the 8oxoG:A Hoogsteen base pairing at SHL_+2_ shows a different structural signature with rapid exchanges between two states (Fig. 8a). The two states are characterized by distinct values of the buckle, propeller, inclination, and tip angles: 130°/−115°/70°/−50° respectively for the first state, and −130°/115°/−50°/60° respectively for the second state (Fig. 8b). The presence of two distinct states is consistent across non-nucleosomal DNA and nucleosomal DNA at SHL_+2_, indicating that the nucleosome does not alter the dynamic nature of the 8oxoG:A base pair. We do however observe subtle differences in the distribution of the two conformational states between non-nucleosomal DNA and nucleosomal DNA at SHL_+2_. These differences could be attributed to the environment of the base pair, namely the local DNA sequence and/or the exposure to interactions with histone tails in the nucleosome (Fig. 4h). The Hoogsteen hydrogen bonds between the nucleobases are stable along the simulations and this dynamical behavior remains very local. Perturbations on the adjacent base pairs are limited to a bimodal distribution of only the 5′ base pair tip angle (SFig. 17). Additionally, in agreement with the cryo-EM structures, the global nucleosome dynamics are not impacted by the presence of the 8oxoG:A base pair. However, we do observe local perturbations of the double-helix structure at the 8oxoG:A base pair at SHL_+2_. The distance between N2 of 8oxoG and the closest non-bridging oxygen at SHL_+2_ is greater than that for H8 of dG and the closest non-bridging oxygen in the ND-NCP system (Fig. 4b and 8c), indicating that there are local rearrangements of the DNA backbone due to the presence of 8oxoG. The Hoogsteen base pairing features the O8 atom of 8oxoG pointing towards the minor groove, near the phosphate groups. This conformation makes it particularly attractive to bridging water molecules and sodium cations. For this reason, water molecules and sodium cations within 3 Å of the O8 of 8oxoG are found in higher quantities in the 8oxoG:A systems as compared to the 8oxoG:C systems (SFig. 8-9) and these interactions can alter the DNA backbone angle distribution. Indeed, a bimodal distribution of the DNA α and γ backbone angles is observed at SHL_+2_, revealing the presence of two different conformations (Fig. 8d). This indicates that the local perturbations of the DNA double-helix structure at the 8oxoG:A base pair at SHL+2 are much more pronounced than for the 8oxoG:C Watson-Crick base pair. Hoogsteen base pairing promotes spatial and orientational changes to nucleobases which disrupt base stacking. This, along with the N2 of 8oxoG amino group pointing towards the major groove might contribute to the local destabilization we observe. These rearrangements of the DNA backbone are observed in both the 8oxoG:A-NCP_+2_ and duplex DNA system (Fig. 8c-d), indicating that the presence of the nucleosome does not affect the 8oxoG:A induced perturbation of the local DNA backbone.

**Fig. 8:**
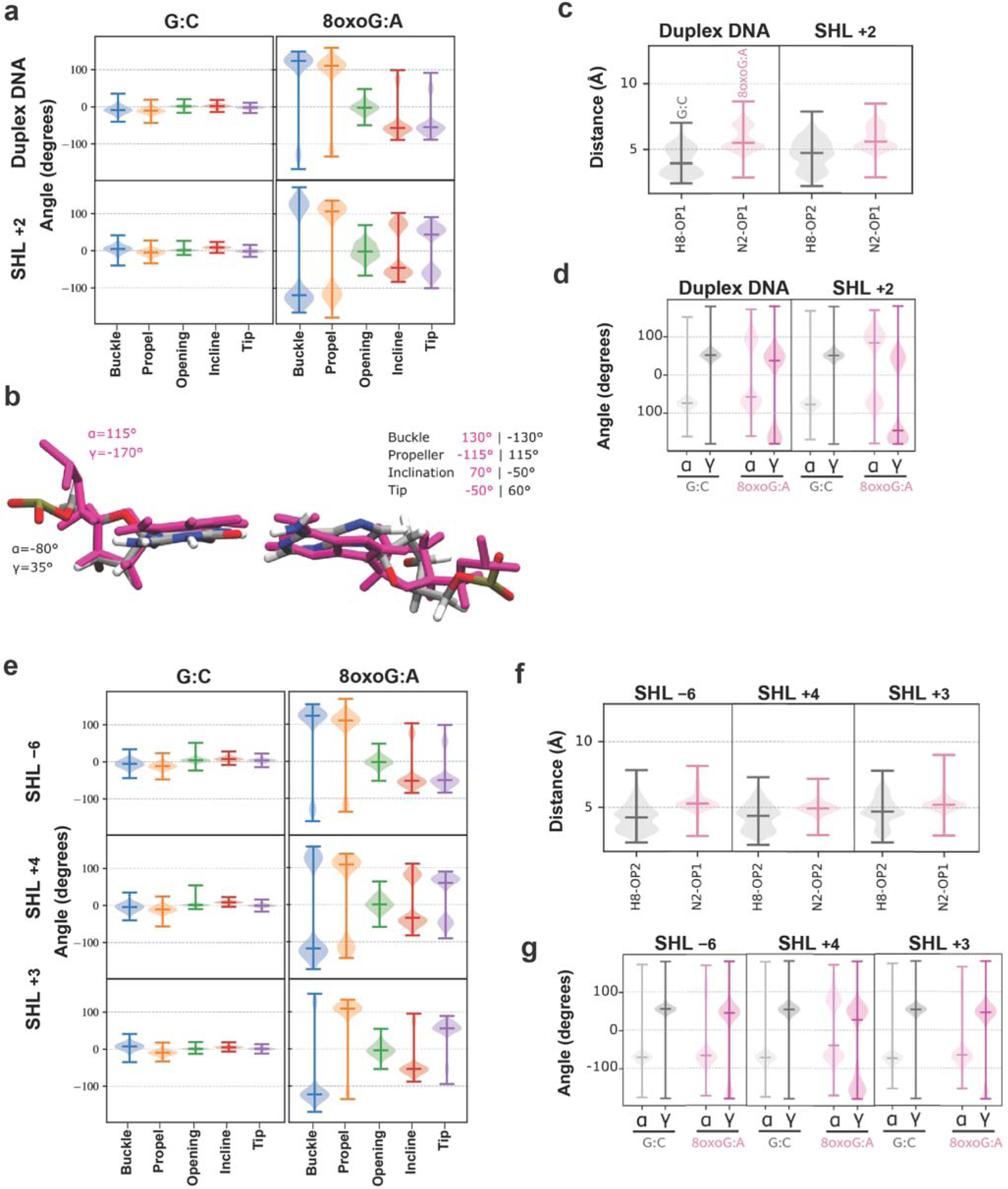
Dynamics of the 8oxoG:A base pair in the nucleosome and non-nucleosomal (duplex) DNA. **a,** Distribution of the intra-base pair angles for the control and 8oxoG:A systems in duplex DNA or at SHL_+2_. **b,** Superimposition of representative snapshots from the simulation with 8oxoG:A., illustrating the two conformations sampled for the 8oxoG:A mismatch. The conformations feature distinct backbone angles and intra-base pair parameter values, some of which are displayed. Structures have been aligned on the nucleotides sugar rings to show the backbone and nucleobase dynamics. **c,** Distribution of the difference between either the H8 (undamaged, grey) or N2 (for 8oxoG:A, pink) atom and the closest non-bridging oxygen atom on the phosphate group (OP1 or OP2) calculated from MD simulations in duplex DNA or SHL_+2_. **d,** Distribution of the α and γ backbone angles of 8oxoG or dG in the control, for duplex DNA or SHL_+2_. **e,** Distribution of the intra-base pair angles for the control and 8oxoG:A systems at SHL_−6_, SHL_+4_, and SHL_+3_. **f,** Distribution of the difference between either the H8 (undamaged, grey) or N2 (for 8oxoG:A, pink) atom and the closest non-bridging oxygen atom on the phosphate group (OP1 or OP2) calculated from MD simulations at SHL_−6_, SHL_+4_, and SHL_+3_. **g,** Distribution of the α and γ backbone angles of 8oxoG or dG in the control, for SHL_−6_, SHL_+4_, and SHL_+3_.

To determine whether the intra-base pair structural parameters of 8oxoG:A and the 8oxoG:A-induced local rearrangements in the DNA backbone are position dependent in the nucleosome, we performed five replicates of 1 µs classical MD simulations on the 8oxoG:A-NCP_+3_, 8oxoG:A-NCP_+4_, and 8oxoG:A-NCP_−6_ structures. We found that at each additional SHL tested, the intra-base pair structural parameters show bimodal distributions like we saw in the 8oxoG:A-NCP_+2_ system (Fig. 8e). This indicates that the dynamical nature of the 8oxoG:A base pair is conserved throughout several positions in the nucleosome. We do however observe subtle position-dependent differences in the distribution of the two conformational states which could be attributed to the different level of exposure to interactions with histone tails and/or the local DNA sequence (Fig. 4h). To determine if the 8oxoG:A-induced local DNA backbone rearrangements are position dependent, we compared the DNA backbone perturbation at SHL_+3_, SHL_+4_, and SHL_−6_ to SHL_+2_. We found that at each position, there are similar perturbations of the double-helix structure at the 8oxoG:A base pair which are more pronounced compared to the 8oxoG:C base pair. In each 8oxoG:A-NCP system, the distance between N2 of 8oxoG and the closest non-bridging oxygen show higher values as compared to the distance between the H8 of dG and the closest non-bridging oxygen in the ND-NCP system (Fig. 8f). We also observed a bimodal distribution of the α and γ backbone angles in all systems containing 8oxoG:A, similar to what we observed in the 8oxoG:A-NCP_+2_ system (Fig. 8g). This bimodality is especially pronounced at SHL_+4_, similar to what we observed at this location with the 8oxoG:C base pair (Fig. 4g). Cumulatively, these data indicate that the 8oxoG:A base pair induces local destabilization of the DNA backbone at each position tested in the nucleosome, though to slightly different extents.

### MUTYH is not appreciably active in the nucleosome

We and others have shown that OGG1 is able to excise solvent-exposed 8oxoG from a nucleosome when the lesion is base paired with dC^27,28,34,35,43,44^. However, whether MUTYH can excise dA paired with 8oxoG at any location in the nucleosome remains unknown. MUTYH achieves substrate specificity and performs excision activity on duplex DNA by making extensive contacts with both DNA strands. Specifically, MUTYH extrudes dA from the helix and into its catalytic pocket while simultaneously rotating the opposing 8oxoG to the *anti* conformation by making contacts with both its C- and N-terminal domains^40^. This engagement of MUTYH with both strands of the duplex elicits a conformational change into a catalytically competent state^45,46^. We suspect the histone core and nucleosomal DNA may hinder the ability of MUTYH to form the interactions necessary for excision of dA when the 8oxoG:A base pair is present in the nucleosome (SFig. 18). Additionally, our 8oxoG:A-NCP structures show that the 8oxoG:A base pair does not cause distortion in the DNA that may facilitate recognition and excision by MUTYH.

To test MUTYH excision activity in a nucleosome, we performed biochemical experiments on a population of nucleosomes where each nucleosome contains at most one 8oxoG:A base pair. Collectively, the population includes nucleosomes with 8oxoG:A at 32 unique locations that span all SHLs. The dA nucleobase is always in the same strand of DNA but has varied levels of solvent exposure (Figure 9a, STable 3). Of note, the 8oxoG:A base pair locations include SHL_+2_, SHL_+3_, and SHL_+4_ which are consistent with the locations in our 8oxoG:A-NCP_+2_, 8oxoG:A-NCP_+3_, and 8oxoG:A-NCP_+4_ structures, respectively. The population of nucleosomes were incubated with excess MUTYH under single-turnover conditions. Any abasic sites generated by excision of dA by MUTYH were converted to strand breaks by treatment with NaOH and visualized via denaturing PAGE (Figure 9b,c)^28,47^. The same duplex DNA used to assemble nucleosomes, but in this case not bound to histones, was used as a positive control for MUTYH activity.

**Fig. 9:**
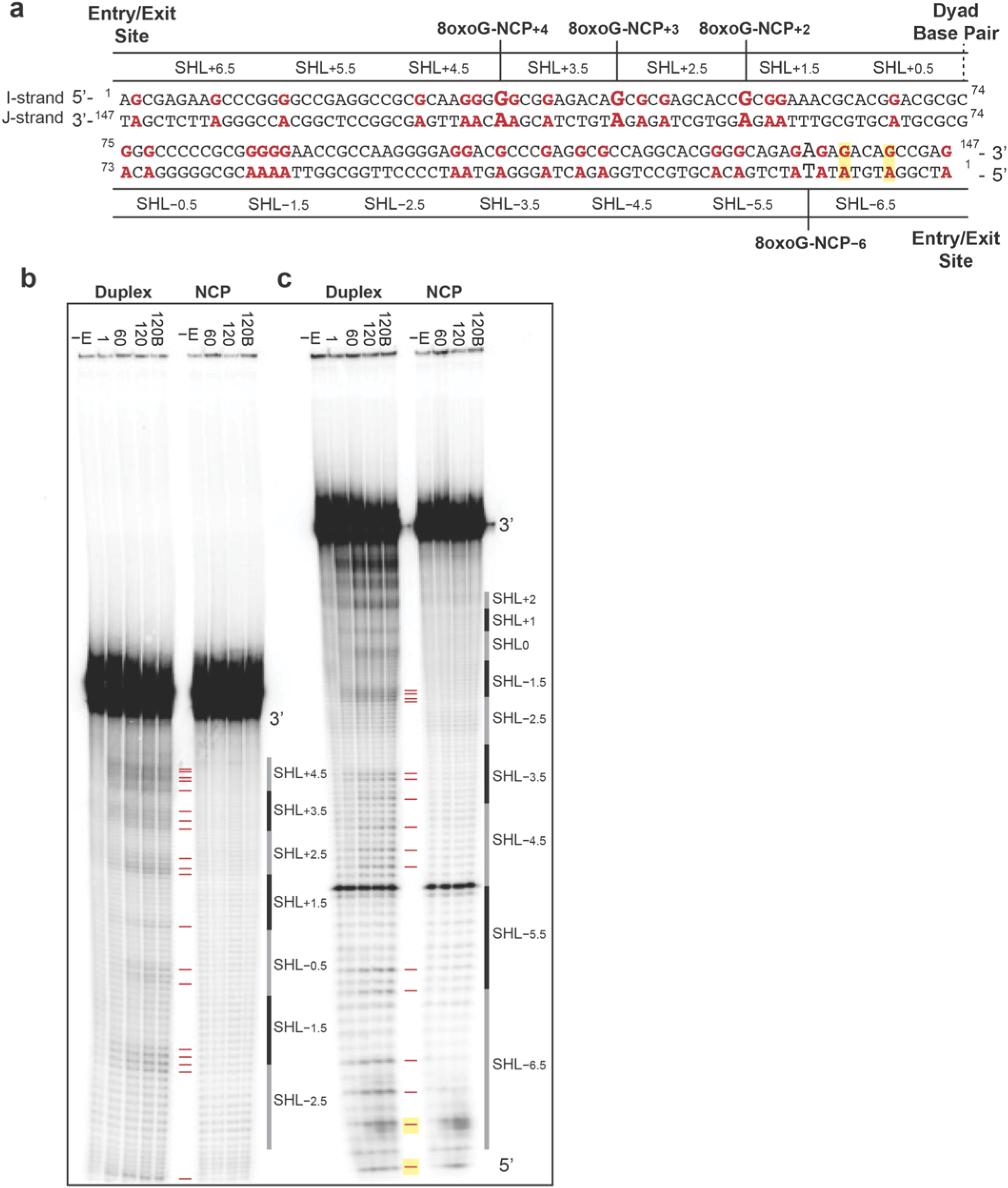
MUTYH can excise dA from 8oxoG:A base pairs in non-nucleosomal DNA, but not nucleosomal DNA. **a,** Diagram of the Widom 601 strong positioning sequence highlighting the globally incorporated 8oxoG:A base pairs. G shown in red represent 8oxoG. Every possible 8oxoG:A location is shown in red. Each I strand contains at most one 8oxoG. **b,** Representative 10 % denaturing PAGE showing dA excision from 8oxoG:A in non-nucleosomal (duplex) and nucleosomal DNA (NCP) by MUTYH. −E samples are substrate (duplex or NCP) incubated for 120 min without MUTYH to reveal background damage that was pre-existing or occurred through sample workup. Substrates (duplex or NCP) incubated with MUTYH are labeled by their incubation time: 1, 60, or 120 min. The 120B sample was incubated with MUTYH for 60 min, supplemented with additional MUTYH, and incubated for an additional 60 min. Red dashes mark locations of dA. The asterisk indicates a loading standard. SHL of the NCP are indicated. Gel was run for 3 h to visualize SHL_−2_ to SHL_+4_. **c,** Same as (b) except the gel was run for 1.5 h to visualize SHL_−6.5_ to SHL_−3_. dA that were excised from NCPs by MUTYH are highlighted in yellow.

Our results reveal MUTYH excises dA from each of the 32 locations in the non-nucleosomal DNA (Figure 9b,c; duplex, red dashes). In contrast, in nucleosomal DNA, the band density at all but two dA locations are comparable to the (−E) negative controls, indicating that MUTYH does not excise dA in the vast majority of the nucleosome, despite there being solvent-exposed dA. MUTYH activity is observed at two sites in the nucleosome, both of which are located at SHL_−6.5_ (Figure 9c; NCP, yellow highlights). At SHL_−6.5_, the DNA is in the entry/exit region of the nucleosome and is susceptible to transient and spontaneous unwrapping^48^. We envision that the increased accessibility of DNA in the entry/exit region likely enables MUTYH to form the extensive contacts with both strands of DNA required for catalysis. Notably, dA excision activity by MUTYH is not observed at SHL_+2_ (8oxoG:A-NCP_+2_), SHL_+3_ (8oxoG:A-NCP_+3_) or SHL_+4_ (8oxoG:A-NCP_+4_). Although the population of NCPs did not contain the 8oxoG:A base pair captured in the 8oxoG:A-NCP_−6_ structure, MUTYH activity was not observed at adjacent locations within SHL_−6_.

## Discussion

Oxidative damage of the nucleobase guanine gives rise to the mutagenic DNA lesion 8oxoG, which occurs across the genome in chromatinized and non-chromatinized DNA^21^. How 8oxoG is accommodated in the nucleosome and how oxidative DNA glycosylases repair 8oxoG lesions in chromatin remains poorly understood^49^. Using a combination of structural biology, classical MD simulations, and enzyme kinetics, we characterized the structure and dynamics of the non-mutagenic 8oxoG:C and mutagenic 8oxoG:A base pairs in the nucleosome. In addition, we determined that MUTYH has no appreciable activity to repair the 8oxoG:A lesion in the nucleosome. Together, these findings enhance our understanding of how oxidative DNA damage is accommodated in nucleosomes and how this damage may lead to mutagenic transversions if left unrepaired.

Prior work established that in non-nucleosomal DNA, 8oxoG adopts the *anti* conformation and utilizes Watson-Crick base pairing opposite dC, whereas 8oxoG adopts the *syn* conformation and utilizes Hoogsteen base pairing opposite dA^40,42^. Recent structures of 8oxoG in the nucleosome revealed that 8oxoG uses Watson-Crick base pairing with dC^36,41^. However, little is known about the base pairing properties of 8oxoG:A in the nucleosome, and whether these properties are position dependent. The cryo-EM structures shown here reveal that nucleosomal 8oxoG will base pair with dC and dA using Watson-Crick and Hoogsteen base pairing, respectively, similar to what was observed in non-nucleosomal DNA. Importantly, these base pairing properties remain nearly identical across the four nucleosome translational positions we tested, indicating that the ability of 8oxoG to adopt *anti* or *syn* conformations is not dependent on the local DNA and/or histone environment. Furthermore, we found that the presence of the 8oxoG lesion in the nucleosome did not cause major perturbations to the histone octamer and induces only very local structural perturbations to the DNA backbone. The MD simulations reveal that in the local DNA backbone, there is an equilibrium between two backbone conformations for 8oxoG:C and 8oxoG:A in both nucleosomal and non-nucleosomal DNA. Additionally, there is specific dynamical behavior of the nucleobases in the 8oxoG:A base pair that was not observed for the 8oxoG:C base pair. Together, this indicates that 8oxoG is readily accommodated in the nucleosome, though the 8oxoG:A base pair is more dynamic compared to the 8oxoG:C base pair. We believe the ability of the nucleosome to accommodate 8oxoG suggests that the presence of 8oxoG also likely has minimal impact on nucleosome assembly and/or disassembly.

The primary mechanism for repairing 8oxoG lesions within chromatinized and non-chromatinized DNA is the BER pathway. The method by which BER is initiated to repair 8oxoG is dependent upon its base pairing partner^7,11–13^, where repair of 8oxoG:C is initiated by OGG1, and repair of 8oxoG:A is initiated by MUTYH. Prior work identified that OGG1 excises 8oxoG in the nucleosome, though with reduced efficiency compared to non-nucleosomal DNA, suggesting that 8oxoG:C can be repaired in the context of chromatin^27,28,33–35^. The reduced efficiency of OGG1 is dictated by the position of the 8oxoG:C in the nucleosome, where solution accessible 8oxoG is excised more readily than histone-occluded 8oxoG^28^. In contrast to OGG1, our data indicates that MUTYH has minimal activity in the nucleosome regardless of the rotational orientation of 8oxoG, though some residual activity was seen at the terminal entry/exit site DNA. Interestingly, our attempts to determine structures of MUTYH bound to nucleosomes containing 8oxoG:A at multiple different positions were unsuccessful (see methods). Because of this, we suspect that MUTYH is unable to engage the 8oxoG:A base pair and/or adopt a catalytically competent state as previously observed in crystal structures of MUTYH on non-nucleosomal DNA containing 8oxoG:A^50^. Furthermore, we believe the residual activity of MUTYH at the nucleosome entry/exit site could be attributed to the dynamic nature of the DNA at that position^27^. When the DNA temporarily dissociates from the histone octamer, MUTYH may encompass the site of DNA damage in a catalytically competent state and successfully excise dA.

Failure to repair an 8oxoG:A base pair will result in a G to T transversion, a common mutation signature in cancer^9,10^. Prior work showed that G to T transversions occur at a higher rate in histone bound DNA compared to linker DNA between nucleosomes following oxidative stress in human cells^21^. We envision that OGG1 can excise 8oxoG in non-nucleosomal DNA, and to a lesser extent, nucleosomal DNA, but that MUTYH does not efficiently excise dA opposite 8oxoG in nucleosomal DNA. The inability of MUTYH to function at most sites in the nucleosome, and the partial ability of OGG1 to function in the nucleosome provides a strong rationale for the observed mutational spectra. Of note, in vitro activity of BER enzymes does not correlate with the magnitude of mutational frequency, suggesting that additional cellular factors, including chromatin remodelers and BER cofactors, likely help facilitate the repair of oxidative

DNA damage in chromatin^51–56^. Additionally, these experiments took advantage of the 601 strong positioning sequence, but in cells, histones and nucleosomal DNA are not as tightly associated^57^. This less rigid association may result in increased ability of MUTYH to repair an 8oxoG:A base pair within the nucleosome. Furthermore, nucleosome remodeling could promote repositioning of a nucleosomal 8oxoG:A base pair to the linker region between nucleosomes that is more accessible to BER proteins. Several chromatin remodelers have been shown to be epistatic with BER factors including Amplified in Liver Cancer 1 (ALC1)^58^, Chromodomain Helicase DNA-binding protein 6 (CHD6)^59^, and HELLS^60^. Future studies will be required to dissect the interplay between these remodelers and MUTYH-initiated BER. Lastly, other BER cofactors could help promote MUTYH activity *in vivo* that is not observed in our *in vitro* experiments. One cofactor that has been shown to be rapidly recruited to oxidative damage within chromatin and stimulate both OGG1 and MUTYH activity is the DNA damage binding protein, UV-DDB^61,62^. It is possible that the addition of UV-DDB to our experiments would lead to increased MUTYH activity, and future studies will be required to address this hypothesis.

## Methods

### Oligonucleotide synthesis, purification, and preparation

For cryo-EM, all non-damaged oligonucleotides (oligos) were obtained from Integrated DNA Technologies (IDT) and all oligos containing 8oxoG were obtained from TriLink BioTechnologies. Each oligo was resuspended at 1 mM in a buffer containing 10 mM Tris-HCl (pH 7.5) and 1 mM EDTA. Each 8oxoG oligo was mixed with its non-damaged complement (see Supplementary Table 4) at equimolar ratios to a final concentration of 500 µM. The oligos were then annealed using a thermal cycler by heating to 95 °C then cooled to 4 °C at a rate of 0.5 °C min^-1^. The annealed oligos were immediately used for nucleosome reconstitution.

For kinetics studies, all DNA was synthesized using phosphoramidite chemistry on a Mermade4 (BioAutomation) DNA synthesizer. Phosphoramidites were purchased from Glen Research. 147-mer DNA was used for duplex controls and NCP samples (SFig. 19). The complementary single strands are denoted as 601I (“I strand”) and 601J (“J strand”) based nomenclature used in an X-ray crystal structure of an NCP containing this Widom 601 DNA^63^. DNA nucleobases are labeled 1 to 147 in the 5’ to 3’ direction for the J strand. 8oxoG was globally incorporated into the I strand using previously reported methods^47^. The Poisson distribution (λ = 0.355) was used to determine the molar ratio to create a T/8oxoG mixture to ensure that 95 % of DNA contained, at most, one 8oxoG per I strand. Both the I and J strands were cleaved from the solid support and protecting groups were removed by incubation with 30% (v/v) NH_4_OH at 55°C for 19 hours, and the I strand (containing 8oxoG) cleavage and deprotection step was supplemented with 0.25 M β-mercaptoethanol (BME) to prevent oxidation of the lesion. The oligonucleotides were purified by 8 % denaturing polyacrylamide gel electrophoresis (PAGE). The purified J strand (containing the dA in the 8oxoG:A base pairs) was 5’-^32^P-radiolabeled and annealed to the 8oxoG-containing I strand in a buffer containing 10 mM Tris (pH 8) and 50 mM NaCl. A single-stranded 27-mer oligonucleotide was used as an internal standard and loading control (5’-GAT GTA TAT ATC TGA CAC GTG CTG GGA-3’). This internal standard was synthesized and purified as described above.

### Purification of recombinant human histones for cryo-EM

Expression and purification of *H. sapien* histones were performed using well-established methods^64,65^. In brief, the genes encoding histones H2A, H2B, H3.2 (C110A), and H4 were previously cloned into a pet3a expression vector. The expression vectors were then transformed into T7 Express lysY competent cells (histone H2A, H3.2, H4; New England Biolabs) or BL21-CodonPlus competent cells (histone H2B; Agilent). The transformed cells were grown in M9 minimal media supplemented with a 1.0% vitamin solution at 37 °C until an OD_600_ of 0.4 was reached. Protein expression was induced with 0.4 mM IPTG (histone H2A, H2B, and H3) or 0.2 mM IPTG (histone H4) for 4 hours (histone H2A and H2B) or 3 hours (histone H3 and H4) at 37°C. The cells were then harvested by centrifugation and resuspended in a buffer containing 50 mM Tris (pH 7.5), 100 mM NaCl, 5 mM BME, and 1 mM EDTA. The resuspended cells were lysed using sonication and inclusion bodies containing each histone were collected via centrifugation. The histones were extracted from the inclusion bodies under denaturing conditions in a buffer containing 20 mM Tris (pH 7.5), 6 M Guanidinium hydrochloride, and 10 mM dithiothreitol (DTT). The extracted histones were dialyzed into 8 M Urea and purified via subtractive anion-exchange and cation-exchange chromatography via gravity flow. The purified histones were dialyzed into H_2_O, lyophilized, and stored long term at −20 °C.

### Preparation of H2A/H2B Dimers and H3/H4 Tetramers for cryo-EM

H2A/H2B dimer and H3/H4 tetramer were generated and refolded using a previously established method^64,65^. In brief, each purified histone was resuspended in a buffer containing 20 mM Tris (pH 7.5), 6 M Guanidinium hydrochloride, and 10 mM DTT. To generate H2A/H2B dimer, equimolar amounts of histone H2A and H2B were combined, dialyzed three times against a buffer containing 20 mM Tris-HCl (pH 7.5), 2 M NaCl, 1 mM EDTA, and 1 mM DTT, and subsequently purified by gel filtration using a Sephacryl S-200 HR (Cytiva). To generate H3/H4 tetramer, equimolar amounts of histone H3 and H4 were combined and dialyzed three times against a buffer containing 20 mM Tris-HCl (pH 7.5), 2 M NaCl, 1 mM EDTA, and 1 mM DTT and purified by gel filtration using a Sephacryl S-200 HR (Cytiva). The purified H2A/H2B dimer and H3/H4 tetramer were stored long-term at −20 °C in 50% glycerol.

### Nucleosome Reconstitution

For cryo-EM, nucleosomes were reconstituted using a previously established method^64,65^. In brief, the annealed 8oxoG DNA (see Supplementary Table 4), H2A/H2B dimer, and H3/H4 tetramer were combined in a 1:2:1 molar ratio in a buffer containing 20 mM Tris-HCl (pH 7.5), 2 M NaCl, 1 mM EDTA, and 1 mM DTT. The mixture was dialyzed against a no salt buffer stepwise to a final salt concentration of less than 150 mM NaCl over a period of 24 hours. The reconstituted nucleosomes were subject to heat-shock at 55 °C for 30 minutes and subsequently purified via 10% - 40% sucrose gradient ultracentrifugation. The purified nucleosomes were stored short term (< 1 month) at 4 °C.

For kinetics assays, nucleosomes were reconstituted with canonical human histone octamers (The Histone Source at Colorado State University) as previously described^66^. In brief, 1 µM 147-mer duplex and 1 µM histone octamer were combined in a 1:1 DNA:octamer molar ratio in a Slide-a-Lyzer dialysis device (0.1 mL capacity, 3.5 kDa MWCO; Thermo Fisher Scientific). Samples were incubated at 4 °C in a buffer containing 10 mM Tris-HCl (pH 7.5), 2 M NaCl, 1 mM EDTA, 1 mM DTT, and 0.5 mg/mL BSA. NaCl concentration was reduced stepwise at 1 h intervals (1.2, 1.0, 0.6 M) and 3 h interval (0 M) via dialysis. The EDTA concentration was reduced to 0 mM during the final hour of dialysis. NCPs were filtered by centrifugation using a Spin-X Centrifuge Tube filter (0.22 µm, Corning Incorporated) to remove any precipitates. Samples were evaluated by 7 % native PAGE (19:1 acrylamide:bisacrylamide, 4 °C, 2.5 h, 155 V, 0.25X Tris-borate-EDTA) to ensure that no excess duplex DNA was present (SFig. 19).

### Human MUTYH expression and purification

The overexpression and purification of MUTYH was performed as previously described^67^. Briefly, *E. coli* BL21 (DE3) co-transformed with the pRKISC and pKJE7 plasmids were used as the expression host. The pRKISC vector encodes the [4Fe-4S] cluster assembly machinery^68^, while pKJE7 expresses the *dnaK*, *dnaJ*, and *grgE* chaperones^69^. Chemical competence was induced using RbCl to facilitate subsequent transformation with the pET28-MBP-MUTYH construct. Cells were recovered in SOC media for 1 h and plated on LB agar containing kanamycin (50 µg/mL), tetracycline (15 µg/mL), and chloramphenicol (34 µg/mL). Colonies were harvested to inoculate 2L of Terrific Broth supplemented with the same antibiotics in a 4 L flat-bottom flask. Cultures were grown at 37 °C with shaking at 180 rpm until reaching OD_600_ > 1.5 (∼6 h), followed by cooling at 4 °C without shaking for 1 h. Protein expression was then induced with 1 mM IPTG, along with the addition of 0.1 g of ferrous sulfate and 0.1 g of ferric citrate. Expression proceeded at 15 °C for 16-24 h with shaking at 180 rpm. Two hours prior to harvesting, the cultures were supplemented with 0.120 g zinc sulfate heptahydrate. Cells were harvested by centrifugation (6,000 × *g*, 10 min, 4 °C) and stored at –80 °C.

For purification, frozen pellets were thawed and resuspended in lysis buffer (30 mM Tris-HCl [pH 7.5], 1 M NaCl, 20 mM 2-mercaptoethanol, and 10% glycerol) supplemented with 1 mM phenylmethylsulfonyl fluoride (PMSF). Cells were lysed by sonication on ice using 20s on/ 40s off cycles with 50% duty cycle and 0.5 amplitude (Branson Sonifier 250), followed by centrifugation at 15,000 × *g* for 50 min at 4 °C. The clarified supernatant was incubated with 1.5 mL Ni^2+^-NTA resin (Qiagen) for 1 h at 4 °C with rotation, then poured over a PD-10 column (Cytiva) for gravity flow.

The resin was washed with 25 mL lysis buffer and eluted with 10 mL elution buffer (30 mM Tris [pH 7.5], 200 mM NaCl, 30 mM 2-mercaptoethanol, 10% glycerol, and 500 mM imidazole). Prompt removal of imidazole and further purification was achieved via heparin affinity chromatography. The eluted sample was loaded onto a 1 mL Heparin column (Cytiva) pre-equilibrated with 10% buffer B (30 mM Tris [pH 7.5], 1 M NaCl, 10% glycerol, 1 mM DTT). Bound protein was eluted using a 10–100% linear gradient of buffer B over 45 min at 1.5 mL/min on an ÄKTA Pure 25L (Cytiva). Fractions of MBP-MUTYH were quantified by UV-Vis to determine yield (MW = 100,888 g/mol, extinction coefficient = 152,470 M^-1^/cm^-1^) and collected for overnight proteolysis using TEV protease for the isolation of MUTYH from the MBP fusion protein. Finally, MUTYH was purified by a second heparin-affinity chromatography using an identical gradient and buffer systems as the first heparin-affinity purification. Fractions containing pure MUTYH were assessed by SDS-PAGE and concentrated using Amicon centrifugal filters (10,000 MWCO). Protein concentration was determined by UV absorbance at 280 nm using an extinction coefficient of 84,630 M^-1^/cm^-1^. Aliquots were stored at –80 °C for future use.

### Cryo-EM sample and grid preparation

To generate the 8oxoG:C samples for cryo-EM, 5 µM of reconstituted nucleosome core particle (NCP) containing 8oxoG damage across from dC was mixed with 7.5 µM – 10 µM of OGG1 K294Q in a buffer containing 25 mM HEPES (pH 7.1), 25 mM NaCl, 1 mM TCEP and 1 mM EDTA and incubated on ice for 10 minutes. The OGG1-NCP complexes were crosslinked in a final concentration of 0.1% glutaraldehyde and incubated on ice for 20 minutes. Directly after incubation, the samples were loaded onto a Superdex 200 Increase 10/300 GL (Cytiva) that had been pre-equilibrated in a buffer containing 50 mM HEPES (pH 7.1), 100mM NaCl, 1mM TCEP, and 1 mM EDTA. The fractions containing NCP were identified via native polyacrylamide gel electrophoresis (5%, 59:1 acrylamide:bis-acrylamide). The samples were concentrated to 1.5 µM for short-term storage. Gels for each 8oxoG:C-NCP sample used for cryo-EM grid preparation can be found in Supplemental Figure 2.

To generate the 8oxoG:A samples for cryo-EM, 12 µM of NCP containing 8oxoG damage across from dA was mixed with 24 µM of MUTYH in a buffer containing 25 mM HEPES (pH 7.5), 50 mM NaCl, 0.5 mM TCEP, and 2.5 mM EDTA and incubated on ice for 10 minutes. The MUTYH-NCP complexes were crosslinked in a final concentration of 0.15% glutaraldehyde and incubated on ice for 20 minutes. Directly after incubation, the samples were loaded onto a Superdex S200 Increase 10/300 GL (Cytiva) that had been pre-equilibrated in a buffer containing 25 mM HEPES (pH 7.5), 50 mM NaCl, 0.5 mM TCEP, and 2.5 mM EDTA. The fractions containing NCP were identified via native polyacrylamide gel electrophoresis (5%, 59:1 acrylamide:bis-acrylamide). The samples were concentrated to 1.5 µM (8oxoG:A-NCP_−6_ and 8oxoG:A-NCP_+4_) or 2 µM (8oxoG:A-NCP_+3_ and 8oxoG:A-NCP_+2_). Gels for each 8oxoG:A-NCP sample used for cryo-EM grid preparation can be found in Supplemental Figures 4, 7, and 10. 3 μL of each concentrated sample for both 8oxoG:C-NCP and 8oxoG:A-NCP samples was applied to a Quantifoil R2/2 300 mesh copper cryo-EM grid and plunge frozen in liquid ethane using a Vitrobot Mark IV (Thermo Fisher). The cryo-EM grids were clipped and stored in liquid nitrogen prior to screening and data collection.

### Cryo-EM Data collection and processing

The cryo-EM data collections for 8oxoG:C-NCP_+3_ and 8oxoG:C-NCP_+2_ were performed using a Titan Krios G3i with Gatan K3 camera and BioQuantum energy filter at the University of Chicago Advanced Electron Microscopy Core Facility with a raw pixel size of 0.534 Å/pixel. The cryo-EM data collections for 8oxoG:A-NCP_−6_, 8oxoG:A-NCP_+4_, 8oxoG:A-NCP_+3_, and 8oxoG:A-NCP_+2_ were performed using a Titan Krios with Falcon 4i camera and Selectris X energy filter at Pacific Northwest Cryo-EM Center with a raw pixel size of 0.394 Å/pixel (8oxoG:A-NCP_−6_, 8oxoG:A-NCP_+4_, 8oxoG:A-NCP_+3_) or 0.4125 Å/pixel (8oxoG:A-NCP_+2_). The cryo-EM data collection for ND-NCP was performed using a Thermo Glacios with Falcon 4i camera and Selectris energy filter at University of Kansas Medical Center Cryo-EM Research Facility with a raw pixel size of 1.18 Å/pixel. The cryo-EM datasets were each processed using cryoSPARC v4^70,71^ and the complete processing workflow for each dataset is shown in Supplemental Figures 2, 4, 7, and 10. In brief, the micrographs were corrected for particle motion resulting from stage drift and beam irradiation using Patch Motion Correction and subsequently contrast transfer function (CTF) fit using Patch CTF. The micrographs were then manually curated to exclude micrographs based on factors including CTF fit resolution and ice thickness. A random subset of the curated micrographs were then used to generate initial 2D classes via blob picker, and these initial 2D classes were used for automated template picking of the complete dataset. The template picked particles were curated and extracted from the micrographs at a box size of 600 pixels and down sampled to 256 pixels. The extracted particles were subjected to multiple rounds of 2D classification to yield a final particle stack. The final particle stacks were used to generate an ab-initio model followed by multiple rounds of heterogenous refinement to separate out destroyed or unwrapped nucleosomes. For all datasets apart from 8oxoG:A-NCP_+2_, 3D classification was performed to improve the interpretability of the map using a focus mask for the entry/exit site DNA, which is often found unwrapped from the histone core. The final particles were re-extracted at a full box size of 600 pixels and subjected to local CTF refinement and final non-uniform refinement. The final maps were deposited into the Electron Microscopy Data Bank (EMDB) under accession numbers EMD-47136 for 8oxoG:C-NCP_+2_, EMD-47137 for ND-NCP, EMD-47138 for 8oxoG:C-NCP_+3_, EMD-47139 for 8oxoG:A-NCP_+2_, EMD-47140 for 8oxoG:A-NCP_+3_, EMD-47141 for 8oxoG:A-NCP_+4_, and EMD-47142 for 8oxoG:A-NCP_−6_.

### Model building and refinement

Model building and refinement for each structure was performed using University of California San Francisco (UCSF) ChimeraX, PHENIX, and COOT^72–76^. A cryo-EM nucleosome structure containing an AP-site at SHL_−6_ (PDB: 7U51)^25^ was used as a starting model for each structure and 8oxoG and adenine (if applicable) were subsequently built in at the locations of damage. The models for each structure were docked into its respective cryo-EM map using ChimeraX. The models were then iteratively refined in PHENIX using secondary structure restraints and manual adjustments were performed to the model using COOT. The final models were validated using MolProbity^77^ and deposited into the Protein Data Bank (PDB) under accession numbers 9DS4 for 8oxoG:C-NCP_+2_, 9DS5 for ND-NCP, 9DS6 for 8oxoG:C-NCP_+3_, 9DS7 for 8oxoG:A-NCP_+2_, 9DS8 for 8oxoG:A-NCP_+3_, 9DS9 for 8oxoG:A-NCP_+4_, and 9SDA for 8oxoG:A-NCP_−6_. All depictions of the maps and models presented in the manuscript were generated using UCSF ChimeraX.

### Molecular Dynamics simulations

Cryo-EM structures were used as starting structure for molecular dynamics simulations of 8oxoG:C-NCP_+2_, 8oxoG:C-NCP_+3_, 8oxoG:A-NCP_−6,_ 8oxoG:A-NCP_+4_, 8oxoG:A-NCP_+3_, and 8oxoG:A-NCP_+2_. In addition, 8oxoG:C-NCP_+4_, 8oxoG:C-NCP_−6_ and control ND-NCP structures were computationally created starting from 8oxoG:C-NCP_+2_ structure. A 21-bp non-nucleosomal, double-stranded DNA sequence was built as the starting model for a control system with the sequence 5′-CGG CTG TAT AG*A TCT GAC AGC-3′ and its complementary sequence (*indicates site of 8oxoG in damaged system). Histone tails were added, or in an extended conformation, or after alignment of the nucleosome core particle with the 1KX5 PDB structure^78^. Five starting conformations of the tails were thus designed per sequence. Each system was solvated in a truncated octahedron box of TIP3P water^79^ with a buffer of 15 Å around the solute. Sodium cations and chloride anions were added to reach neutrality and a physiological saline concentration of 0.10 M.

The MD simulations were performed using Amber20 facilities^80^. Protein and DNA were described using ff14SB^81^ and parmbsc1 force field^82^ force fields respectively. 8oxoG force field was taken from literature^83^. Protonation states have been determined according to PropKa calculations^84^. CUFIX correction was applied to improve ionic interactions^85^. Hydrogen mass repartitioning (HMR) method^86^ were used to allow a timestep of 4 fs for all equilibration and production trajectories. All simulations were run in periodic boundary conditions with a cut-off of 10 Å for non-covalent interactions and Particle-Mesh Ewald approach for electrostatic interaction. First, all systems were minimized with 5,000 steps of steepest descent followed by 5,000 steps using conjugate gradient. Then, they were heated from 0 K to 300 K during 30 ps, with a timestep of 1 fs, using a Langevin thermostat with a collision frequency of 1 ps^-1^ in the NVT ensemble. An equilibration run at 300 K was performed during 100 ns, with a pressure of 1 atm using the Berendsen barostat. Finally, a production of 1 μs was performed for each system, totaling 5 replicas per sequence. The same protocol was applied to model the 8oxoG:C and 8oxo:A in duplex DNA. Starting structure consisting in a 21 base pairs oligonucleotide centered on the lesion were extracted from the cryo-EM structures featuring the lesion at SHL_−_6, and 3 replicas of 1 µs were collected for each system (8oxoG:C, 8oxoG:A and undamaged control). All MD data is available online on a Zenodo repository (10.5281/zenodo.14755037).

Base pair parameters were computed using the courbes program^87^. This Python wrapper uses Curves+^88^ as a backend and automates the extraction of critical descriptors from multiple replicas of molecular dynamics simulations, generating basic statistical reports and facilitating efficient data analysis. Courbes is freely available at https://github.com/rglez/courbes. The radial distributions of water molecules and sodium ions were computed with the cpptraj program of Amber20.

### MUTYH excision activity assays

To assess MUTYH activity, 50 nM of either duplex or NCPs was incubated at 37 °C with 500 nM active MUTYH in 20 mM Tris (pH = 7.6), 50 mM NaCl, 150 mM KCl, 1 mM DTT, 0.2 mg/mL BSA in a total reaction volume of 82 µL. At various time intervals, a 20 µL aliquot was removed and quenched with 20 µL of 1 M NaOH for 2 min at 90 °C. The time intervals used were 1, 60, and 120 min for duplex and 60 and 120 min for NCPs. A fresh aliquot of MUTYH (10 pmol) was added to the 120B sample after 60 min of incubation at 37°C. A negative-control sample (−E) was incubated in the absence of MUTYH at 37 °C and underwent the same quench and workup as other samples to reveal MUTYH-independent strand cleavage. DNA fragments were extracted from proteins with 25:24:1 phenol:chloroform:isoamyl alcohol and desalted by ethanol precipitation. Samples were resuspended in 50 % aqueous formamide and loaded onto a 10 % denaturing PAGE. Half of each sample was subject to electrophoresis for 3 h to resolve SHL_−1.5_ to SHL_+4.5_, and the other half was subject to electrophoresis for 1.5 h to resolve SHL_−6.5_ to SHL_−2.5_. Gels were imaged by phosphoimagery.

## Acknowledgements

This research was supported by the National Institutes of General Medical Science R35GM128562 (B.D.F.), National Science Foundation MCB-2111680 (S.D.), and the National Cancer Institute R01CA067985 (S.S.D.). J.C.F. has been supported by a training grant from the National Institute of Environmental Health Sciences (T32ES007272). A portion of this research was supported by NIH grant R24GM154185 and performed at the Pacific Northwest Center for Cryo-EM (PNCC) with assistance from Marzia Miletto and Nancy Meyer. A portion of this research was supported by S10OD036339 and performed at the University of Kansas Medical Center Electron Microscopy Research Laboratory with assistance from David Ingham. We thank Tyler M. Weaver for help training A.F.V. and J.A.L. in cryo-EM data processing, model building, and data analysis. We acknowledge Dana Biechele-Speziale for conducting preliminary biochemical experiments.

## Contributions

A.F.V., J.A.L., and B.D.F. conceptualized the experiments and established research goals. J.A.L, J.C.F., and C.S.J. generated nucleosomes for biochemical and cryo-EM experiments. J.A.L performed cryo-EM sample preparation and validation. A.F.V. and J.A.L. processed and analyzed cryo-EM datasets. A.F.V. performed model building and refinement. N.G., R.G., Y.Q., and E.B. performed and analyzed classical molecular dynamic simulations. M.H., C.H.T., and S.D. generated purified MUTYH for biochemical experiments. J.C.F., C.S.J., and S.D. performed and analyzed biochemical experiments. A.F.V., J.A.L., and B.D.F. wrote the manuscript with assistance from J.C.F., C.S.J., E.B., S.S.D, and S.D.

