## Supplemental Material for "Structure, dynamics, and processing of 8oxoG:A in the nucleosome"

for

**Table of Contents**

**Figure S1.** Nucleosome reconstitution for cryo-EM studies………………..……………..….……S3

**Figure S2.** SPA processing workflow for 8oxoG:C-NCP+2….…………………………………..…S4

**Figure S3.** 8oxoG:C-NCP+2 map and model quality assessment….…………………………..S5-6

**Figure S4.** SPA processing workflow for ND-NCP…………………………………………………S7

**Figure S5.** ND-NCP map and model quality assessment….……………………………………S8-9

**Figure S6.** SPA processing workflow for 8oxoG:C-NCP+3…………………………………..……S10

**Figure S7.** 8oxoG:C-NCP+3 map and model quality assessment……………………….……S11-12

**Figure S8.** Solvation shell around the 8oxoG O8 atom….………………………………….…S13-14

**Figure S9.** Na^+^ cations distribution around the 8oxoG O8 atom………………………...……S15-16

**Figure S10.** SPA processing workflow for 8oxoG:A-NCP+2….………………………………...…S17

**Figure S11.** 8oxoG:A-NCP+2 map and model quality assessment….…………………….…S18-19

**Figure S12.** SPA processing workflow for 8oxoG:A+3………………………………………..……S20

**Figure S13.** 8oxoG:A-NCP+3 map and model quality assessment….…………………….…S21-22

**Figure S14.** SPA processing workflow for 8oxoG:A-NCP+4 and 8oxoG:A-NCP−6…………S23-24

**Figure S15.** 8oxoG:A-NCP+4 map and model quality assessment….…………………….…S25-26

**Figure S16.** 8oxoG:A-NCP−6 map and model quality assessment….…………………….…S27-28

**Figure S17.** Intra-base pair parameters 5′ of the damage site….…………………….………S29-30

**Figure S18.** Modeled MUTYH recognition mechanism in the NCP….……………………………S31

**Figure S19.** Nucleosome assembly for enzyme kinetics studies…………………………..……S32

**Table S1.** Cryo-EM table for ND-NCP and 8oxoG:C NCPs......................................................S33

**Table S2.** Cryo-EM table for 8oxoG:A NCPs............................................................................S34

**Table S3.** Solvent Exposure and MUTYH Excision Activity on dA Sites..................................S35

**Table S4.** DNA oligonucleotides used for reconstitution of NCPs for cryo-EM studies…...S36-37

**Supplementary Fig. 1**

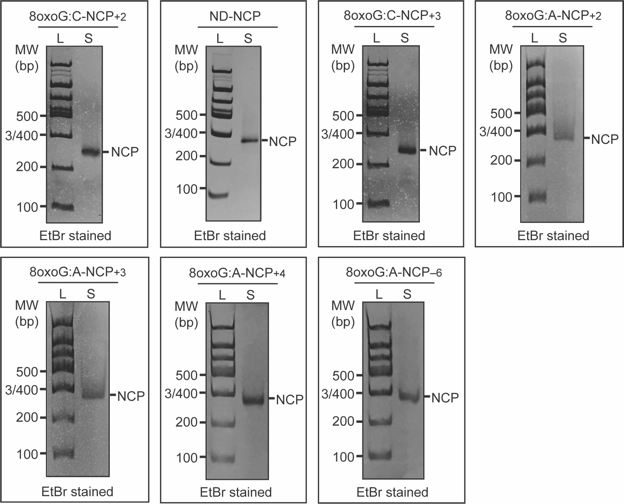

**Supplementary Fig. 1: Nucleosome reconstitution for cryo-EM studies**

Native PAGE gels confirming nucleosome formation and purity for the 8oxoG:C-NCP+2, ND-NCP, 8oxoG:C-NCP+3, 8oxoG:A-NCP+2, 8oxoG:A-NCP+4, and 8oxoG:A-NCP–6. The native PAGE gels were run immediately after initial nucleosome formation and purification. The NCPs were detected using ethidium bromide staining. The 100 bp DNA ladder (L) and nucleosome sample (S) are labeled.

**Supplementary Fig. 2**

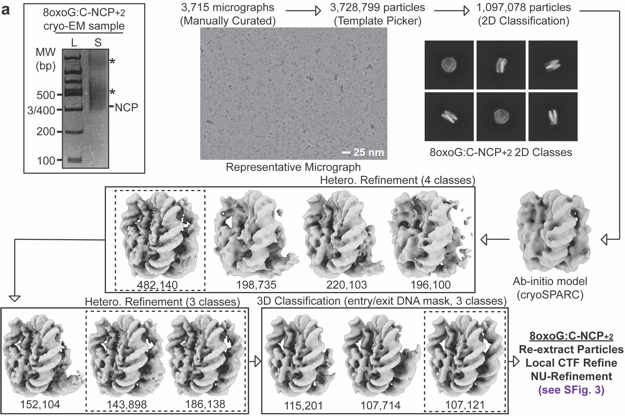

**Supplementary Fig. 2: SPA processing workflow for 8oxoG:C-NCP+2**

**a,** Native PAGE gel of 8oxoG:C-NCP+2 cryo-EM sample and flowchart of the data processing pipeline for the 8oxoG:C-NCP+2 cryo-EM dataset. A 100 bp DNA ladder (L) and the cryo-EM sample (S) are labeled and the NCP was detected using ethidium bromide staining. * indicates higher MW contaminants. A representative micrograph and representative 2D classes from the 8oxoG:C-NCP+2 cryo-EM dataset are shown. The final maps, final models, and quality assessment metrics for 8oxoG:C-NCP+2 can be found in Supplementary Fig. 3.

**Supplementary Fig. 3**

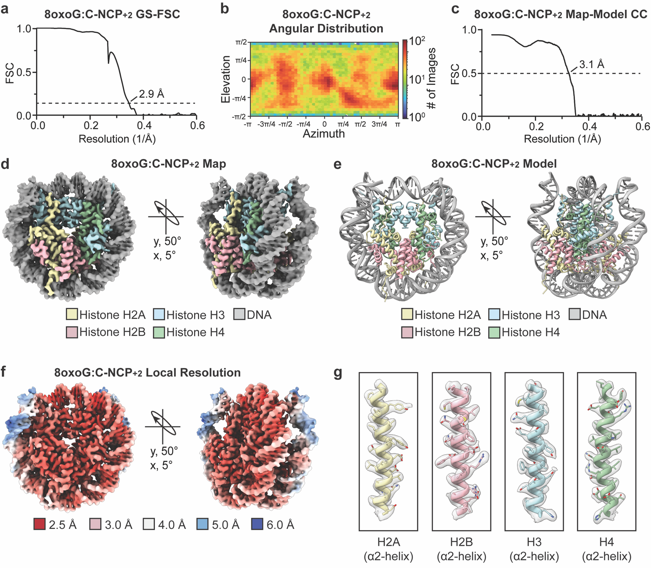

**Supplementary Fig. 3: 8oxoG:C-NCP+2 map and model quality assessment**

**a,** Gold-standard Fourier shell correlation (GS-FSC) curve for the 8oxoG:C-NCP+2 cryo-EM map (black solid line). The dashed line corresponds to FSC – 0.143. **b,** Angular distribution heatmap for the 8oxoG:C-NCP+2 cryo-EM map. **c,** Map-to-model FSC curve for the 8oxoG:C-NCP+2 model and 8oxoG:C-NCP+2 cryo-EM map. The dashed line corresponds to FSC – 0.5. **d,** The final 2.9 Å 8oxoG:C-NCP+2 cryo-EM map shown in two orientations. **e,** The final 8oxoG:C-NCP+2 model shown in two orientations. **f,** The local resolution estimation for the 8oxoG:C-NCP+2 cryo-EM map shown in two orientations. **g,** Representative segmented densities for histones H2A, H2B, H3, and H4 in the 8oxoG:C-NCP+2 cryo-EM map. The representative segmented densities from the cryo-EM map are shown as transparent gray surfaces.

**Supplementary Fig. 4**

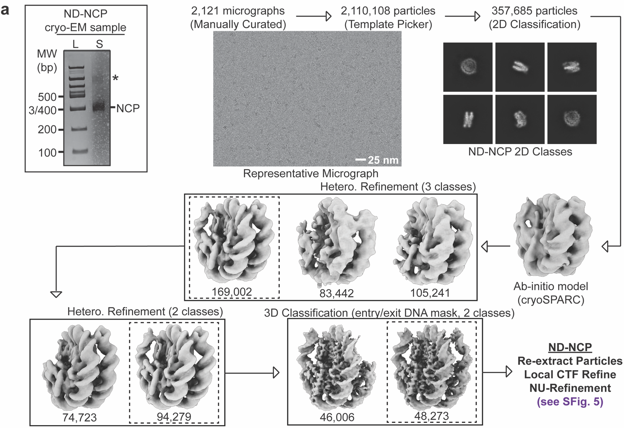

**Supplementary Fig. 4: SPA processing workflow for ND-NCP**

**a,** Native PAGE gel of ND-NCP cryo-EM sample and flowchart of the data processing pipeline for the ND-NCP cryo-EM dataset. A 100 bp DNA ladder (L) and the cryo-EM sample (S) are labeled and the NCP was detected using ethidium bromide staining. * indicates higher MW contaminants. A representative micrograph and representative 2D classes from the ND-NCP cryo-EM dataset are shown. The final maps, final models, and quality assessment metrics for ND-NCP can be found in Supplementary Fig. 5.

**Supplementary Fig. 5**

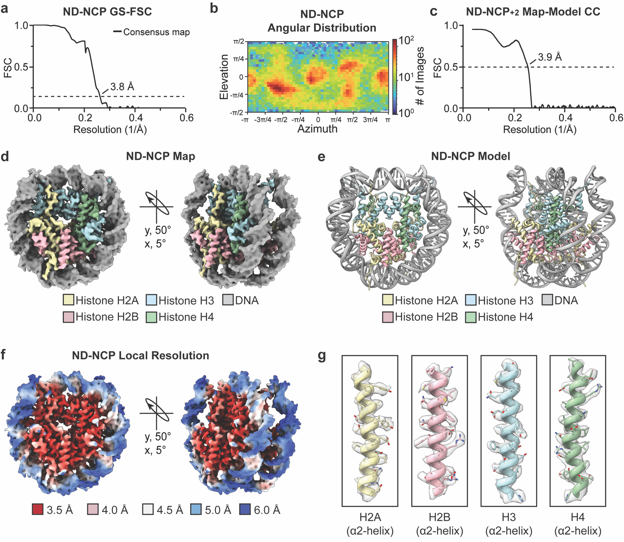

**Supplementary Fig. 5: ND-NCP map and model quality assessment**

**a,** Gold-standard Fourier shell correlation (GS-FSC) curve for the ND-NCP cryo-EM map (black solid line). The dashed line corresponds to FSC – 0.143. **b,** Angular distribution heatmap for the ND-NCP cryo-EM map. **c,** Map-to-model FSC curve for the ND-NCP model and ND-NCP cryo-EM map. The dashed line corresponds to FSC – 0.5. **d,** The final 3.8 Å ND-NCP cryo-EM map shown in two orientations. **e,** The final ND-NCP model shown in two orientations. **f,** The local resolution estimation for the ND-NCP cryo-EM map shown in two orientations. **g,** Representative segmented densities for histones H2A, H2B, H3, and H4 in the ND-NCP cryo-EM map. The representative segmented densities from the cryo-EM map are shown as transparent gray surfaces.

**Supplementary Fig. 6**

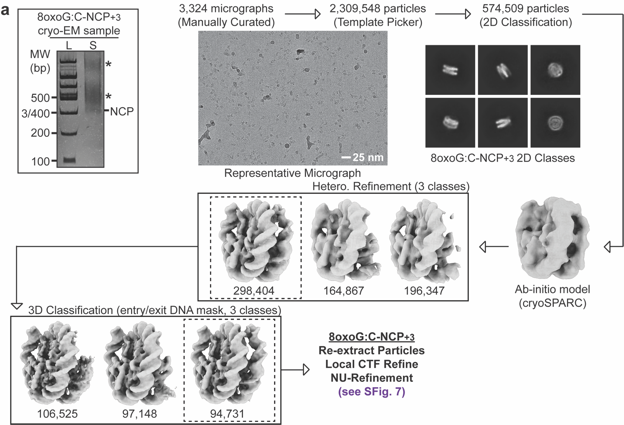

**Supplementary Fig. 6: SPA processing workflow for 8oxoG:C-NCP+3**

**a,** Native PAGE gel of 8oxoG:C-NCP+3 cryo-EM sample and flowchart of the data processing pipeline for the 8oxoG:C-NCP+3 cryo-EM dataset. A 100 bp DNA ladder (L) and the cryo-EM sample (S) are labeled and the NCP was detected using ethidium bromide staining. * indicates higher MW contaminants. A representative micrograph and representative 2D classes from the 8oxoG:C-NCP+3 cryo-EM dataset are shown. The final maps, final models, and quality assessment metrics for 8oxoG:C-NCP+3 can be found in Supplementary Fig. 7.

**Supplementary Fig. 7**

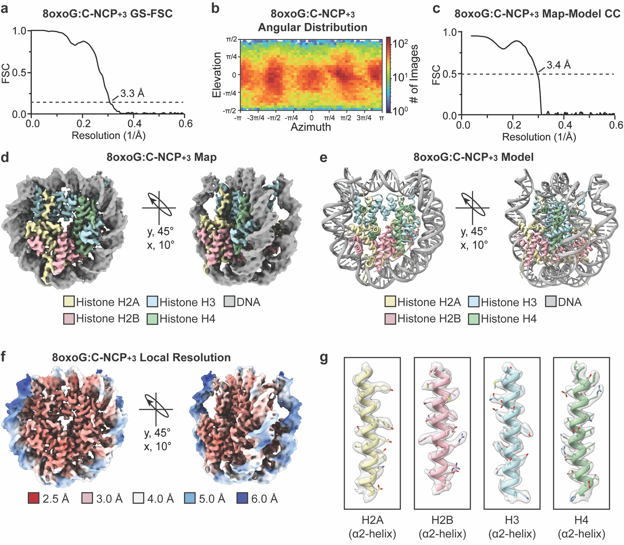

**Supplementary Fig. 7: 8oxoG:C-NCP+3 map and model quality assessment**

**a,** Gold-standard Fourier shell correlation (GS-FSC) curve for the 8oxoG:C-NCP+3 cryo-EM map (black solid line). The dashed line corresponds to FSC – 0.143. **b,** Angular distribution heatmap for the 8oxoG:C-NCP+3 cryo-EM map. **c,** Map-to-model FSC curve for the 8oxoG:C-NCP+3 model and 8oxoG:C-NCP+3 cryo-EM map. The dashed line corresponds to FSC – 0.5. **d,** The final 3.3 Å 8oxoG:C-NCP+3 cryo-EM map shown in two orientations. **e,** The final 8oxoG:C-NCP+3 model shown in two orientations. **f,** The local resolution estimation for the 8oxoG:C-NCP+3 cryo-EM map shown in two orientations. **g,** Representative segmented densities for histones H2A, H2B, H3, and H4 in the 8oxoG:C-NCP+3 cryo-EM map. The representative segmented densities from the cryo-EM map are shown as transparent gray surfaces.

**Supplementary Fig. 8**

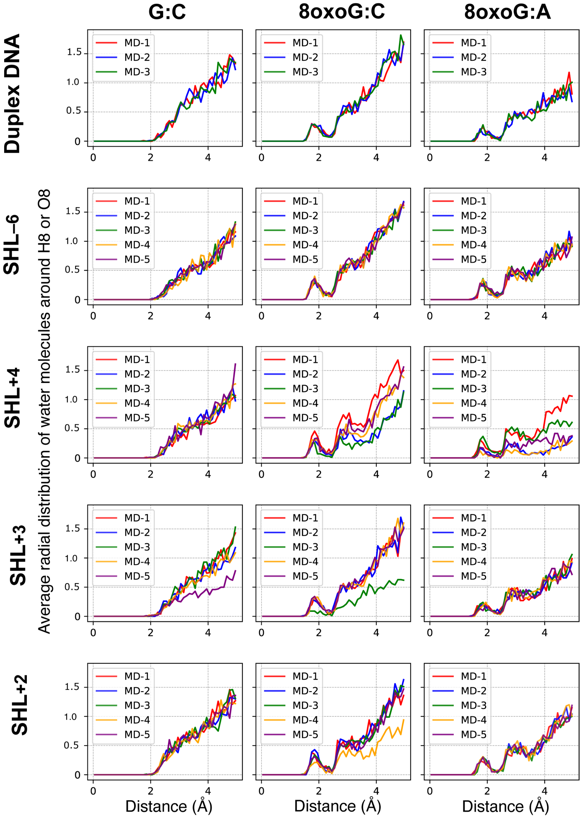

**Supplementary Fig. 8: Solvation shell around the 8oxoG O8 atom**

Average radial distribution of the water molecules around the H8 atom in the control (left), or the O8 atom in the 8oxoG:C (center) and 8oxoG:A (right) systems. A first solvation shell is systematically observed around O8.

**Supplementary Fig. 9**

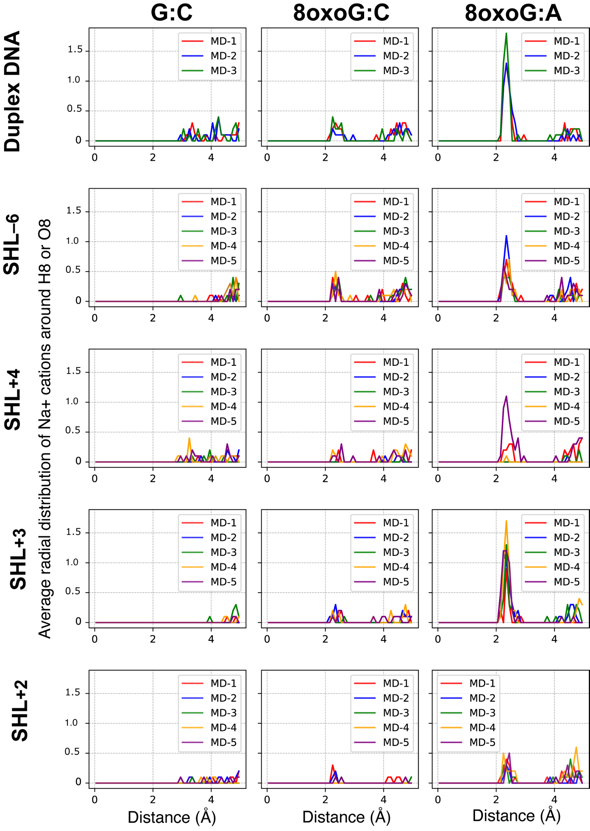

**Supplementary Fig. 9: Na^+^ cations distribution around the 8oxoG O8 atom**

Average radial distribution of the sodium cations around the H8 atom in the control (left), or the O8 atom in the 8oxoG:C (center) and 8oxoG:A (right) systems. The Hoogsteen pairing favors the presence of Na^+^ around O8.

**Supplementary Fig. 10**

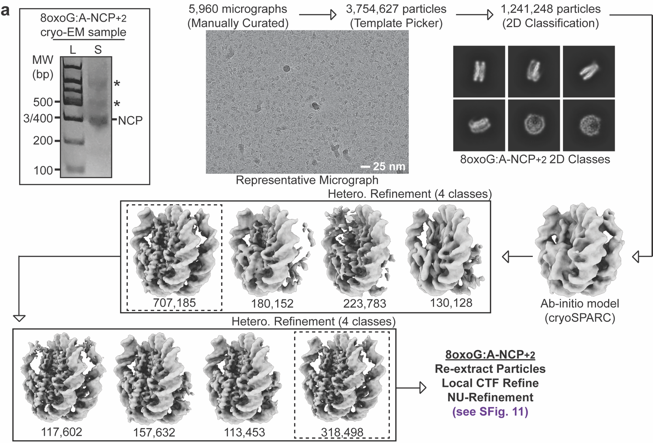

**Supplementary Fig. 10: SPA processing workflow for 8oxoG:A-NCP+2**

**a,** Native PAGE gel of 8oxoG:A-NCP+2 cryo-EM sample and flowchart of the data processing pipeline for the 8oxoG:A-NCP+2 cryo-EM dataset. A 100 bp DNA ladder (L) and the cryo-EM sample (S) are labeled and the NCP was detected using ethidium bromide staining. * indicates higher MW contaminants. A representative micrograph and representative 2D classes from the 8oxoG:A-NCP+2 cryo-EM dataset are shown. The final maps, final models, and quality assessment metrics for 8oxoG:A-NCP+2 can be found in Supplementary Fig. 11.

**Supplementary Fig. 11**

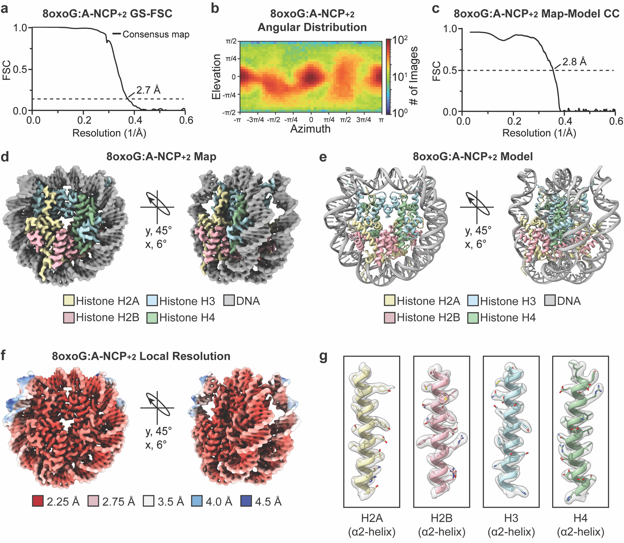

**Supplementary Fig. 11: 8oxoG:A-NCP+2 map and model quality assessment**

**a,** Gold-standard Fourier shell correlation (GS-FSC) curve for the 8oxoG:A-NCP+2 cryo-EM map (black solid line). The dashed line corresponds to FSC – 0.143. **b,** Angular distribution heatmap for the 8oxoG:A-NCP+2 cryo-EM map. **c,** Map-to-model FSC curve for the 8oxoG:A-NCP+2 model and 8oxoG:A-NCP+2 cryo-EM map. The dashed line corresponds to FSC – 0.5. **d,** The final 2.7 Å 8oxoG:A-NCP+2 cryo-EM map shown in two orientations. **e,** The final 8oxoG:A-NCP+2 model shown in two orientations. **f,** The local resolution estimation for the 8oxoG:A-NCP+2 cryo-EM map shown in two orientations. **g,** Representative segmented densities for histones H2A, H2B, H3, and H4 in the 8oxoG:A-NCP+2 cryo-EM map. The representative segmented densities from the cryo-EM map are shown as transparent gray surfaces.

**Supplementary Fig. 12**

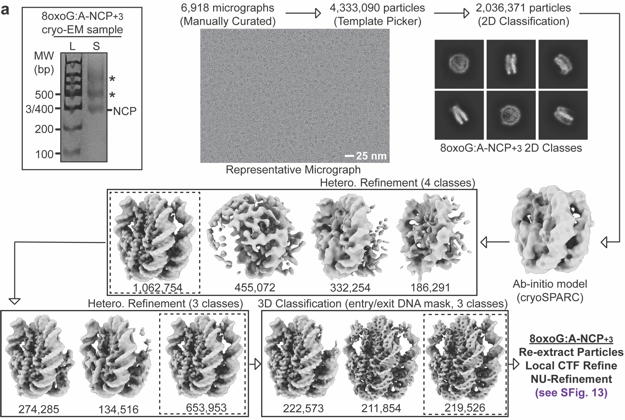

**Supplementary Fig. 12: SPA processing workflow for 8oxoG:A-NCP+3**

**b,** Native PAGE gel of 8oxoG:A-NCP+3 cryo-EM sample and flowchart of the data processing pipeline for the 8oxoG:A-NCP+3 cryo-EM dataset. A 100 bp DNA ladder (L) and the cryo-EM sample (S) are labeled and the NCP was detected using ethidium bromide staining. * indicates higher MW contaminants. A representative micrograph and representative 2D classes from the 8oxoG:A-NCP+3 cryo-EM dataset are shown. The final maps, final models, and quality assessment metrics for 8oxoG:A-NCP+3 can be found in Supplementary Fig. 13.

**Supplementary Fig. 13**

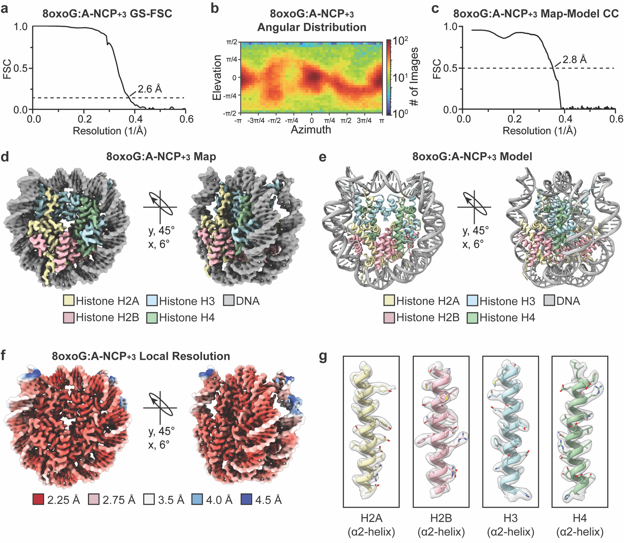

**Supplementary Fig. 13: 8oxoG:A-NCP+3 map and model quality assessment**

**a,** Gold-standard Fourier shell correlation (GS-FSC) curve for the 8oxoG:A-NCP+3 cryo-EM map (black solid line). The dashed line corresponds to FSC – 0.143. **b,** Angular distribution heatmap for the 8oxoG:A-NCP+3 cryo-EM map. **c,** Map-to-model FSC curve for the 8oxoG:A-NCP+3 model and 8oxoG:A-NCP+3 cryo-EM map. The dashed line corresponds to FSC – 0.5. **d,** The final 2.6 Å 8oxoG:A-NCP+3 cryo-EM map shown in two orientations. **e,** The final 8oxoG:A-NCP+3 model shown in two orientations. **f,** The local resolution estimation for the 8oxoG:A-NCP+3 cryo-EM map shown in two orientations. **g,** Representative segmented densities for histones H2A, H2B, H3, and H4 in the 8oxoG:A-NCP+3 cryo-EM map. The representative segmented densities from the cryo-EM map are shown as transparent gray surfaces.

**Supplementary Fig. 14**

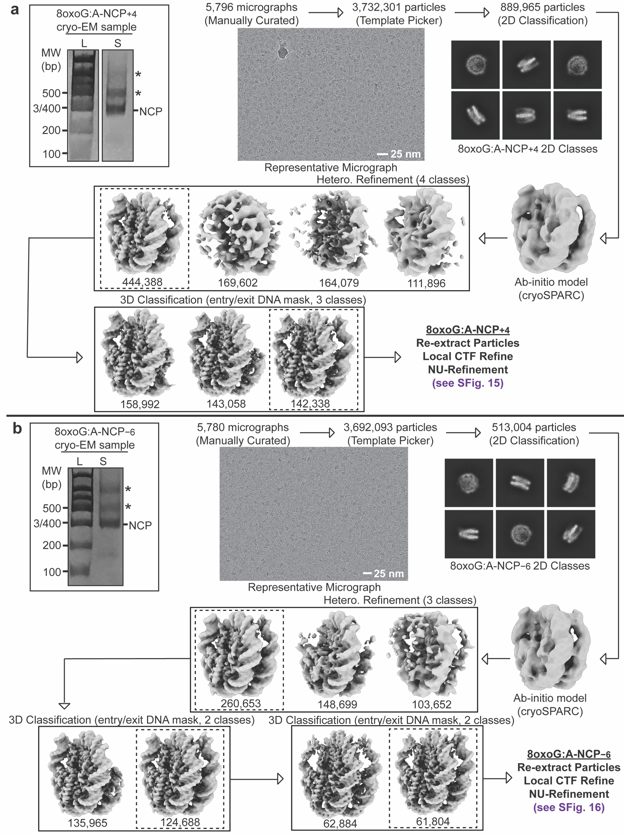

**Supplementary Fig. 14: SPA processing workflow for 8oxoG:A-NCP+4 and 8oxoG:A-NCP−6**

**a,** Native PAGE gel of 8oxoG:A-NCP+4 cryo-EM sample and flowchart of the data processing pipeline for the 8oxoG:A-NCP+4 cryo-EM dataset. A 100 bp DNA ladder (L) and the cryo-EM sample (S) are labeled and the NCP was detected using ethidium bromide staining. * indicates higher MW contaminants. A representative micrograph and representative 2D classes from the 8oxoG:A-NCP+4 cryo-EM dataset are shown. The final maps, final models, and quality assessment metrics for 8oxoG:A-NCP+4 can be found in Supplementary Fig. 15. **b,** Native PAGE gel of 8oxoG:A-NCP−6 cryo-EM sample and flowchart of the data processing pipeline for the 8oxoG:A-NCP−6 cryo-EM dataset. A 100 bp DNA ladder (L) and the cryo-EM sample (S) are labeled and the NCP was detected using ethidium bromide staining. * indicates higher MW contaminants. A representative micrograph and representative 2D classes from the 8oxoG:A-NCP−6 cryo-EM dataset are shown. The final maps, final models, and quality assessment metrics for 8oxoG:A-NCP−6 can be found in Supplementary Fig. 16.

**Supplementary Fig. 15**

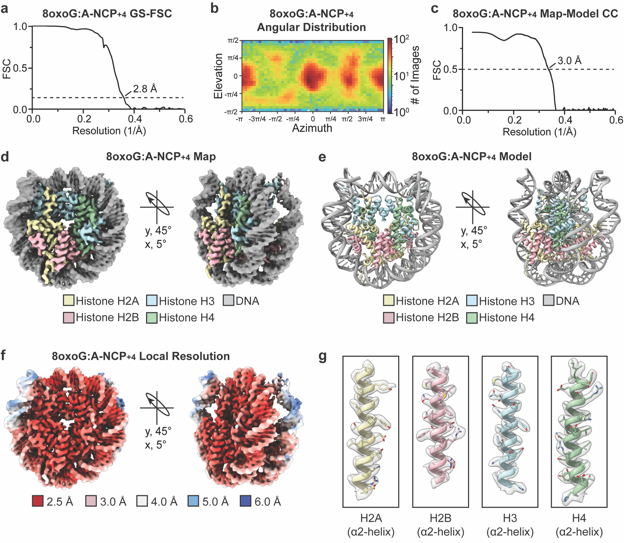

**Supplementary Fig. 15: 8oxoG:A-NCP+4 map and model quality assessment**

**a,** Gold-standard Fourier shell correlation (GS-FSC) curve for the 8oxoG:A-NCP+4 cryo-EM map (black solid line). The dashed line corresponds to FSC – 0.143. **b,** Angular distribution heatmap for the 8oxoG:A-NCP+4 cryo-EM map. **c,** Map-to-model FSC curve for the 8oxoG:A-NCP+4 model and 8oxoG:A-NCP+4 cryo-EM map. The dashed line corresponds to FSC – 0.5. **d,** The final 2.8 Å 8oxoG:A-NCP+4 cryo-EM map shown in two orientations. **e,** The final 8oxoG:A-NCP+4 model shown in two orientations. **f,** The local resolution estimation for the 8oxoG:A-NCP+4 cryo-EM map shown in two orientations. **g,** Representative segmented densities for histones H2A, H2B, H3, and H4 in the 8oxoG:A-NCP+4 cryo-EM map. The representative segmented densities from the cryo-EM map are shown as transparent gray surfaces.

**Supplementary Fig. 16**

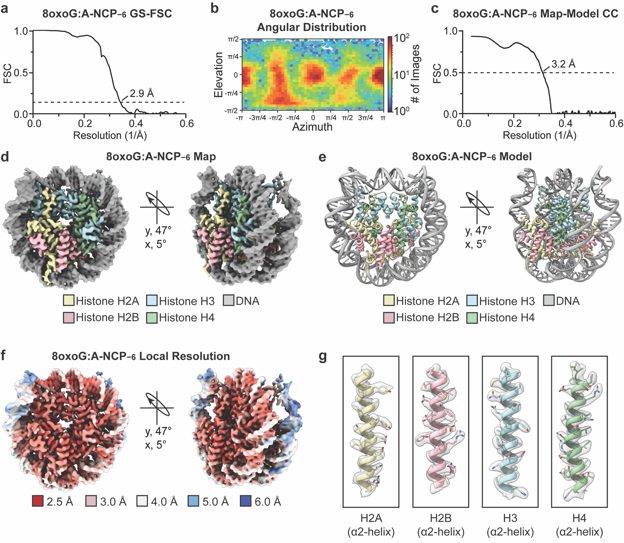

**Supplementary Fig. 16: 8oxoG:A-NCP−6 map and model quality assessment**

**a,** Gold-standard Fourier shell correlation (GS-FSC) curve for the 8oxoG:A-NCP−6 cryo-EM map (black solid line). The dashed line corresponds to FSC – 0.143. **b,** Angular distribution heatmap for the 8oxoG:A-NCP−6 cryo-EM map. **c,** Map-to-model FSC curve for the 8oxoG:A-NCP−6 model and 8oxoG:A-NCP−6 cryo-EM map. The dashed line corresponds to FSC – 0.5. **d,** The final 2.9 Å 8oxoG:A-NCP−6 cryo-EM map shown in two orientations. **e,** The final 8oxoG:A-NCP−6 model shown in two orientations. **f,** The local resolution estimation for the 8oxoG:A-NCP−6 cryo-EM map shown in two orientations. **g,** Representative segmented densities for histones H2A, H2B, H3, and H4 in the 8oxoG:A-NCP−6 cryo-EM map. The representative segmented densities from the cryo-EM map are shown as transparent gray surfaces.

**Supplementary Fig. 17**

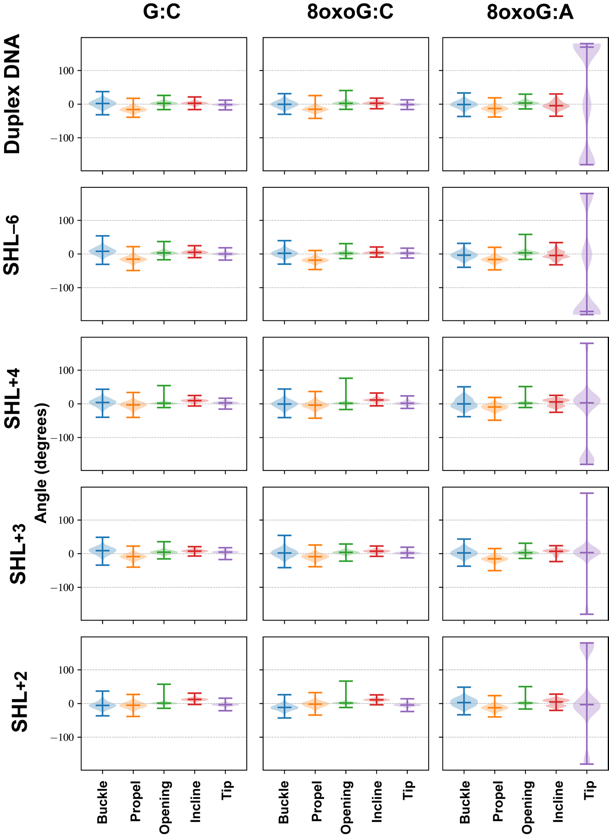

**Supplementary Fig. 17: Intra-base pair parameters 5′ of the damage site**

Distribution of intra-base pair angles of the base pair 5′ of the damage site, for the control (left), the 8oxoG:C (center) and the 8oxoG:A (right) systems, in naked DNA or in the nucleosome at the four different SHL.

**Supplementary Fig. 18**

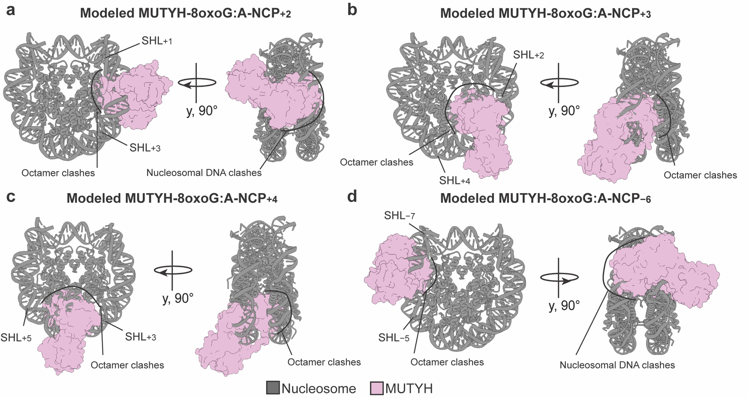

**Supplementary Fig. 18: Modeled MUTYH recognition mechanism in the NCP**

Structural models of MUTYH positioned to interact with 8oxoG:A base pairs at (a) SHL+2, (b) SHL+3, (c) SHL+4, and (d) SHL−6. MUTYH is shown as a surface representation. Significant clashes between MUTYH and the histone octamer and/or nucleosomal DNA are labeled.

**Supplementary Fig. 19**

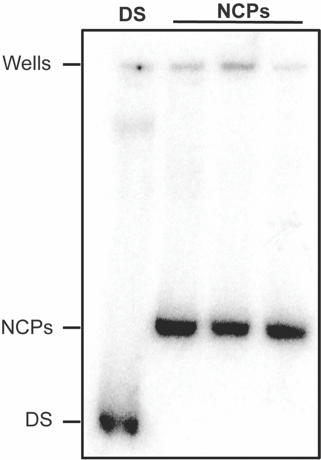

**Supplementary Fig. 19: Nucleosome assembly for enzyme kinetics studies**

Representative native PAGE analyzing NCP assembly. The double stranded DNA control is labeled DS. The three NCP lanes reflect migration of the NCP samples relative to control.

**Supplementary Table 1**

| **Data collection and processing for ND-NCP and 8oxoG:C-NCPs** | | | |
| --- | --- | --- | --- |
| **Dataset** | **ND-NCP** | **8oxoG:C NCP+3** | **8oxoG:C NCP+2** |
| Magnification | 100,000x | 81,000x | 81,000x |
| Voltage (kV) | 200 | 300 | 300 |
| Electron exposure (e^–^/Å^2^) | 40 | 60 | 60 |
| Defocus range (µm) | −0.4 to −1.2 | −0.5 to −2.5 | −0.5 to −2.5 |
| Pixel size (Å) | 1.18 | 0.534 | 0.534 |
| Symmetry imposed | C1 | C1 | C1 |
| Initial particle images (no.) | 2,109,381 | 2,309,548 | 3,728,799 |
| Final particle images (no.) | 48,273 | 94,382 | 106,752 |
| Map resolution (Å) | 3.8 | 3.3 | 2.9 |
| FSC threshold | 0.143 | 0.143 | 0.143 |
| PDB accession | 9DS5 | 9DS6 | 9DS4 |
| EMDB accession | EMD-47137 | EMD-47138 | EMD-47136 |
| **Refinement** | | | |
| Initial model used (PDB ID) | 7U51 | 7U51 | 7U51 |
| Model resolution (Å) | 3.9 | 3.4 | 3.1 |
| FSC threshold | 0.5 | 0.5 | 0.5 |
| **Model composition** | | | |
| Nonhydrogen atoms | 12,030 | 12,044 | 12,052 |
| Protein residues | 758 | 759 | 759 |
| Nucleotide | 294 | 294 | 294 |
| **B factors (Å^2^ )** | | | |
| Protein | 62.06 | 96.98 | 62.61 |
| Nucleotide | 124.71 | 159.56 | 113.75 |
| **r.m.s. deviations** | | | |
| Bond Length (Å) (# > 4σ) | 0.007 (0) | 0.005 (1) | 0.006 (4) |
| Bond Angles (^o^) (# > 4σ) | 1.117 (20) | 0.814 (21) | 0.856 (42) |
| **Validation** | | | |
| MolProbity score | 1.73 | 1.40 | 1.33 |
| Clashscore | 5.95 | 3.83 | 2.90 |
| Poor rotamers (%) | 1.27 | 0.64 | 0.32 |
| **Ramachandran plot** | | | |
| Favored (%) | 95.28 | 96.50 | 96.23 |
| Allowed (%) | 4.72 | 3.50 | 3.77 |
| Disallowed (%) | 0.00 | 0.00 | 0.00 |

**Supplementary Table 2**

| **Data collection and processing for 8oxoG:A-NCPs** | | | |  |
| --- | --- | --- | --- | --- |
| **Dataset** | **8oxoG:A NCP−6** | **8oxoG:A NCP+4** | **8oxoG:A NCP+3** | **8oxoG:A NCP+2** |
| Magnification | 29,000x | 29,000x | 29,000x | 29,000x |
| Voltage (kV) | 300 | 300 | 300 | 300 |
| Electron exposure (e^–^/Å^2^) | 50 | 50 | 50 | 50 |
| Defocus range (µm) | −0.8 to −2.2 | −0.8 to −2.2 | −0.8 to −2.2 | −0.8 to −2.2 |
| Pixel size (Å) | 0.394 | 0.394 | 0.394 | 0.4125 |
| Symmetry imposed | C1 | C1 | C1 | C1 |
| Initial particle images (no.) | 3,692,093 | 3,732,301 | 4,333,090 | 1,096,830 |
| Final particle images (no.) | 61,588 | 141,735 | 218,771 | 316,392 |
| Map resolution (Å) | 2.9 | 2.8 | 2.6 | 2.7 |
| FSC threshold | 0.143 | 0.143 | 0.143 | 0.143 |
| PDB accession | 9DSA | 9DS9 | 9DS8 | 9DS7 |
| EMDB accession | EMD-47142 | EMD-47141 | EMD-47140 | EMD-47139 |
| **Refinement** | | | |  |
| Initial model used (PDB ID) | 7U51 | 7U51 | 7U51 | 7U51 |
| Model resolution (Å) | 3.2 | 3.0 | 2.8 | 2.8 |
| FSC threshold | 0.5 | 0.5 | 0.5 | 0.5 |
| **Model composition** | | | |  |
| Nonhydrogen atoms | 11,991 | 12,039 | 12,107 | 12,094 |
| Protein residues | 753 | 758 | 765 | 762 |
| Nucleotide | 294 | 294 | 294 | 294 |
| **B factors (Å^2^ )** | | | |  |
| Protein | 77.18 | 68.34 | 126.48 | 93.40 |
| Nucleotide | 135.57 | 121.50 | 84.64 | 145.71 |
| **r.m.s. deviations** | | | |  |
| Bond Length (Å) (# > 4σ) | 0.007 (1) | 0.006 (0) | 0.007 (2) | 0.007 (3) |
| Bond Angles (^o^) (# > 4σ) | 1.010 (55) | 0.846 (27) | 0.925 (37) | 0.868 (41) |
| **Validation** | | | |  |
| MolProbity score | 1.75 | 1.31 | 1.70 | 1.46 |
| Clashscore | 5.56 | 3.46 | 4.86 | 4.17 |
| Poor rotamers (%) | 1.28 | 0.79 | 1.73 | 1.10 |
| **Ramachandran plot** | | | |  |
| Favored (%) | 94.57 | 97.04 | 96.13 | 96.51 |
| Allowed (%) | 5.43 | 2.96 | 3.87 | 3.49 |
| Disallowed (%) | 0.00 | 0.00 | 0.00 | 0.00 |

**Supplementary Table 3**

| **SHL of 8oxoG:A** | **Site of dA on J Strand** | **Solvent Exposure of dA** | **Excision of dA from 8oxoG:A by MUTYH** | **Corresponding cryo-EM Structure** |
| --- | --- | --- | --- | --- |
| –6.8 | 6 | MID | Observed | - |
| –6.4 | 10 | LOW | Observed | - |
| –6.2 | 12 | MID | Not Observed | - |
| –6 | 14 | HIGH | Not Observed | - |
| –5.5 | 19 | LOW | Not Observed | - |
| –5.3 | 21 | LOW | Not Observed | - |
| –4.3 | 31 | LOW | Not Observed | - |
| –4.1 | 33 | MID | Not Observed | - |
| –3.8 | 36 | HIGH | Not Observed | - |
| –3.4 | 40 | LOW | Not Observed | - |
| –3.1 | 43 | MID | Not Observed | - |
| –3 | 44 | HIGH | Not Observed | - |
| –1.5 | 59 | MID | Not Observed | - |
| –1.4 | 60 | LOW | Not Observed | - |
| –1.3 | 61 | LOW | Not Observed | - |
| –1.2 | 62 | MID | Not Observed | - |
| –0.3 | 71 | LOW | Not Observed | - |
| –0.1 | 73 | MID | Not Observed | - |
| 0.6 | 80 | LOW | Not Observed | - |
| 1.6 | 90 | LOW | Not Observed | - |
| 1.7 | 91 | LOW | Not Observed | - |
| 1.9 | 93 | MID | Not Observed | 8oxoG:A-NCP+2 |
| 2.6 | 100 | MID | Not Observed | - |
| 2.8 | 102 | LOW | Not Observed | - |
| 3 | 104 | MID | Not Observed | 8oxoG:A-NCP+3 |
| 3.6 | 110 | MID | Not Observed | - |
| 3.9 | 113 | LOW | Not Observed | - |
| 4 | 114 | MID | Not Observed | 8oxoG:A-NCP+4 |
| 4.2 | 116 | HIGH | Not Observed | - |
| 4.3 | 117 | HIGH | Not Observed | - |
| 4.7 | 121 | LOW | Not Observed | - |

**Supplementary Table 3.** Solvent exposure and MUTYH excision activity of dA sites in NCP 8oxoG:dA mispairs. Solvent exposure was assigned based on structural data.

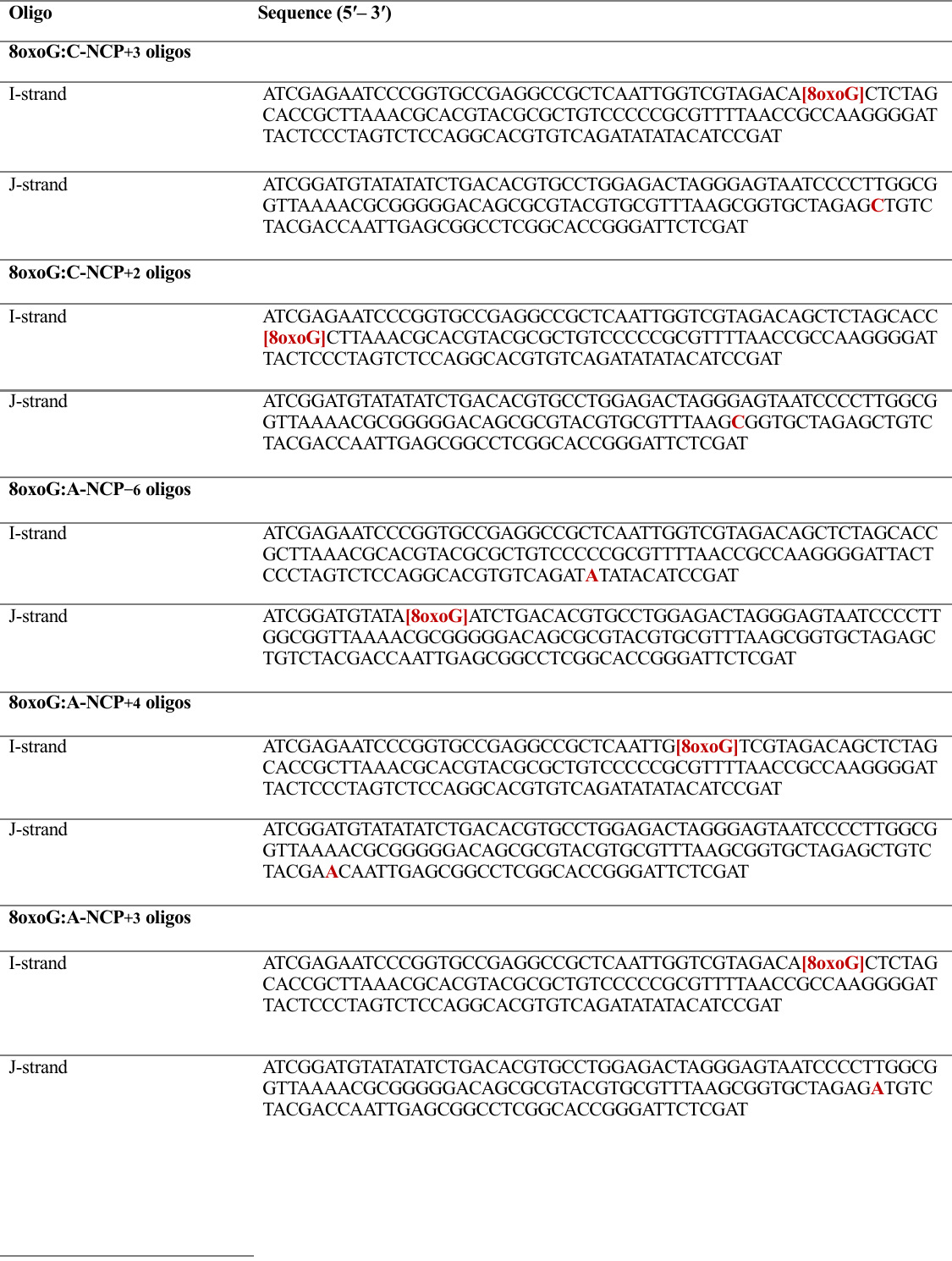
**Supplementary Table 4**

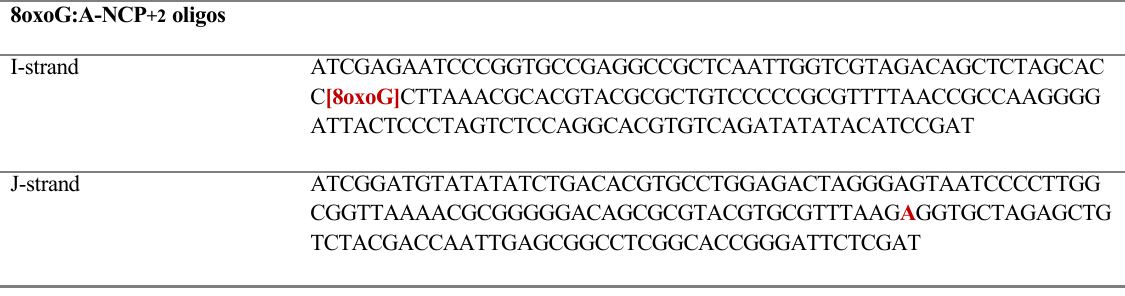

**Supplementary Table 4.** DNA oligonucleotides used for reconstitution of NCPs for cryo-EM studies
